# The ‘Janus A’ gene encodes a polo-kinase whose loss creates a dorsal/ventral intracellular homeosis in the ciliate, *Tetrahymena*

**DOI:** 10.1101/2024.12.19.629484

**Authors:** Eric S. Cole, Wolfgang Maier, Huy Vo Huynh, Benjamin Reister, Deborah Oluwabukola Sowunmi, Uzoamaka Chukka, Chinkyu Lee, Jacek Gaertig

**Author notes:** To whom correspondence should be directed.

## Abstract

Genetic studies on the protist, *Tetrahymena thermophila* provide a glimpse into the unexpectedly rich world of intracellular patterning that unfolds within the ciliate cell cortex. Ciliate pattern studies provide a useful counterpoint to animal models of pattern formation in that the unicellular model draws attention away from fields of cells (or nuclei) as the principal players in the metazoan pattern paradigm, focusing instead on fields of ciliated basal bodies serving as sources of positional information. In this study, we identify *JANA*, a Polo kinase of *Tetrahymena*, that serves as an important factor driving global, circumferential pattern. Loss of function of JanA results in global, mirror-duplication of ventral organelles on the dorsal surface: a kind of intracellular homeosis that has been named the ‘janus’ phenotype. Gain of function (over-expression) reduces or even eliminates cortical organelles within the ventral ‘hemi-cell’. GFP-tagging reveals that JanA decorates basal bodies predominantly within the left-dorsal hemi-cell. These results led us to propose a model in which the default state of cortical patterning is a mirror-image assemblage of cortical organelles including oral apparatus, contractile vacuole pores and cytoproct. JanA normally suppresses organelle assembly in the dorsal hemi-cellular cortex, resulting in a simple, ventral assemblage of these organelles, a ‘half-pattern’ as it were. PLK inhibitors produce a janus phenocopy, but reveal other unanticipated roles for PLK activities involving more local patterning events that control organelle dimensions and organization. We discuss results in light of metazoan studies in which PLK activity links cell cycle control to intracellular symmetry breaking.

## Introduction

### The ciliate cell cortex during pre-division morphogenesis

Ciliated protozoa assemble a suite of complex organelles at precise geometric locations around the cell cortex (Cole & Gaertig, 2022; Cole, et al., 2023). In the decades between 1970 and 2000, a gallery of cortical pattern mutants was established for the model ciliate, *Tetrahymena thermophila* (reviewed by Frankel, 2008; Cole and Gaertig, 2022). Analyses of ciliate pattern phenotypes have revealed pattern mechanisms at work driving anterior-posterior, dorsal-ventral and even left-right (chiral) asymmetry.

Comparative next-generation-sequencing has begun to identify gene products associated with cortical patterning in this ciliate including those involved in anterior-posterior positioning of the fission zone (FZ) and the developing oral primordium (OP) (Fig 1). Surprisingly, the pattern genes discovered so far are often highly conserved proteins including conserved kinases, kinase anchor-proteins (AKAPs), and kinase regulators such as cyclin E and Hippo signaling proteins (Cole, et al., 2023; Tavares, et al., 2012; Jiang, et al., 2017, 2019a, 2020).

**Figure 1.**
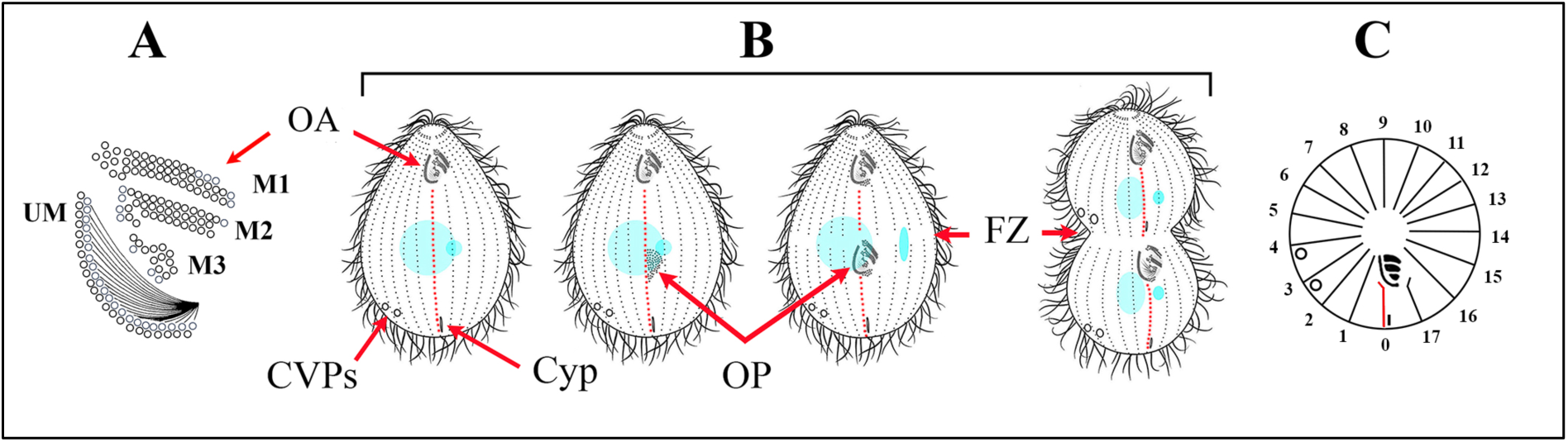
Schematic diagram showing cortical organelles of *Tetrahymena.* A) The oral apparatus (OA) with primary membranelles: M1, M2, M3 and the ‘undulating membrane’ or UM. B) A ventral view of a cell undergoing pre-division morphogenesis showing contractile vacuole pores (CVPs), the cytoproct (Cyp), the developing oral primordium (OP), and the fission zone (FZ). ‘Dots’ represent ciliated basal bodies assembled into ‘kineties’ or ciliary rows. Red dots indicate the ‘stomatogenic kinety’. C) A polar-projection of a non-dividing cell with ciliary rows depicted as lines numbered 0-17. The periphery of the circle represents the posterior end of the cell with the Cyp at kinety 0, CVPs at kineties 3 and 4, and the first post-oral ciliary row (the ‘stomatogenic kinety’) assigned the number 0. The center of the circle represents the anterior end of the cell.

Pattern formation, as studied by most developmental biologists, explores mechanisms that drive differential gene expression within multicellular, multinucleate developmental landscapes. *Tetrahymena* appear to control pattern not through differential gene expression, (there is only one nucleus) but by creating spatial gradients of phosphorylation activity broadcast from fields of ciliated basal bodies. This could rightfully be viewed as a new paradigm of pattern formation.

‘Janus’ mutants represent one of the keystone phenotypes characterized in the Frankel lab (Jerka-Dziadosz & Frankel, 1979, Frankel & Jenkins, 1979). These mutants exhibit a mirror-duplication of the ventral pattern of cortical organelles on the dorsal cell surface. It should be noted, we have adopted the convention of defining the ventral hemi-cell as the region of cortex including ciliary rows 0-4 and 14-17 as in Fig 1C. This region includes all the major cortical organelles. Dorsal, is then assigned to the cortical region from rows 4-14, devoid of organelles other than the ciliary rows themselves. *Janus* mutations, in effect, create a homeotic transformation of the dorsal hemi-cell, into a mirror-image of the ventral hemi-cell.

This mirror pattern duplication is reminiscent of the *bicoid* mutant in *Drosophila*, in which embryos develop a global, mirror-duplication of the posterior half of the embryo within the anterior part of the embryo, or *gooseberry*, in which each segment exhibits a local mirror-duplication of the posterior compartment within the anterior half-segment (Nüsslein-Volhard and Wieschaus, 1980; Frohnhöfer & Nüsslein-Volhard, 1987). Janus mutations in *Tetrahymena* (for which three distinct loci have been identified) orchestrate a global, mirrored pattern-duplication of ventral organelles (oral apparatus, contractile vacuole pores and cytoproct) on the dorsal surface.

### A second-site, janA enhancer mutation (eja)

The original janus mutant (*janA*), was discovered in the very first attempt to isolate temperature-sensitive cell division mutants in *Tetrahymena* (Bruns & Sanford, 1978). From a cell culture mutagenized by exposure to nitrosoguanidine and made homozygous through a form of self-fertilization called ‘short circuit genomic exclusion’ (Bruns, et al., 1976), a clone with enlarged, and somewhat ‘flattened’ features was isolated (CU127). This mutant clone was initially identified due to its high mortality following prolonged growth at elevated temperature (39.5 C). The cell’s cortical features were subsequently analyzed using traditional silver-impregnation staining (Frankel & Heckmann, 1968; Nelsen & DeBault, 1978). CU127 cells exhibited an elevated number of ciliary rows (21-25 in the mutant vs 17-21 seen in a wildtype clone) while 50% of the mutant cells exhibited two sets of CVPs, and 30% exhibited a secondary oral apparatus located 180 degrees around the circumference from the first, ‘normal’ set. This mutant was later named *janA-1* (Jerka-Dziadosz & Frankel, 1979).

Genetic analysis revealed that the *janA*-1 mutation was recessive, mapped to the right arm of chromosome 3, and appeared to require a second-site enhancer mutation (*eja*: enhancer of *janA*) for elevated penetrance (Frankel and Jenkins, 1979). Subsequent rounds of mutant screening recovered a second allele that failed to complement the first (*janA-2*).

### The janus cortical phenotype and its developmental emergence

Taking advantage of *Tetrahymena*’s unique genetics (a somatic, transcriptionally active macronucleus and a transcriptionally silent germline micronucleus), Frankel and Nelsen (1985) were able to observe the progressive emergence of the janus, loss-of-function phenotype within an initially wild-type background. During cell division, the anterior division product of a ciliate (the ‘proter’) inherits the parental OA and assembles a new set of CVPs (nCVPs), while the posterior division product (the ‘opisthe’) inherits the parental CVPs while assembling a new oral primordium (OP) (Figs 2A-F).

**Figure 2.**
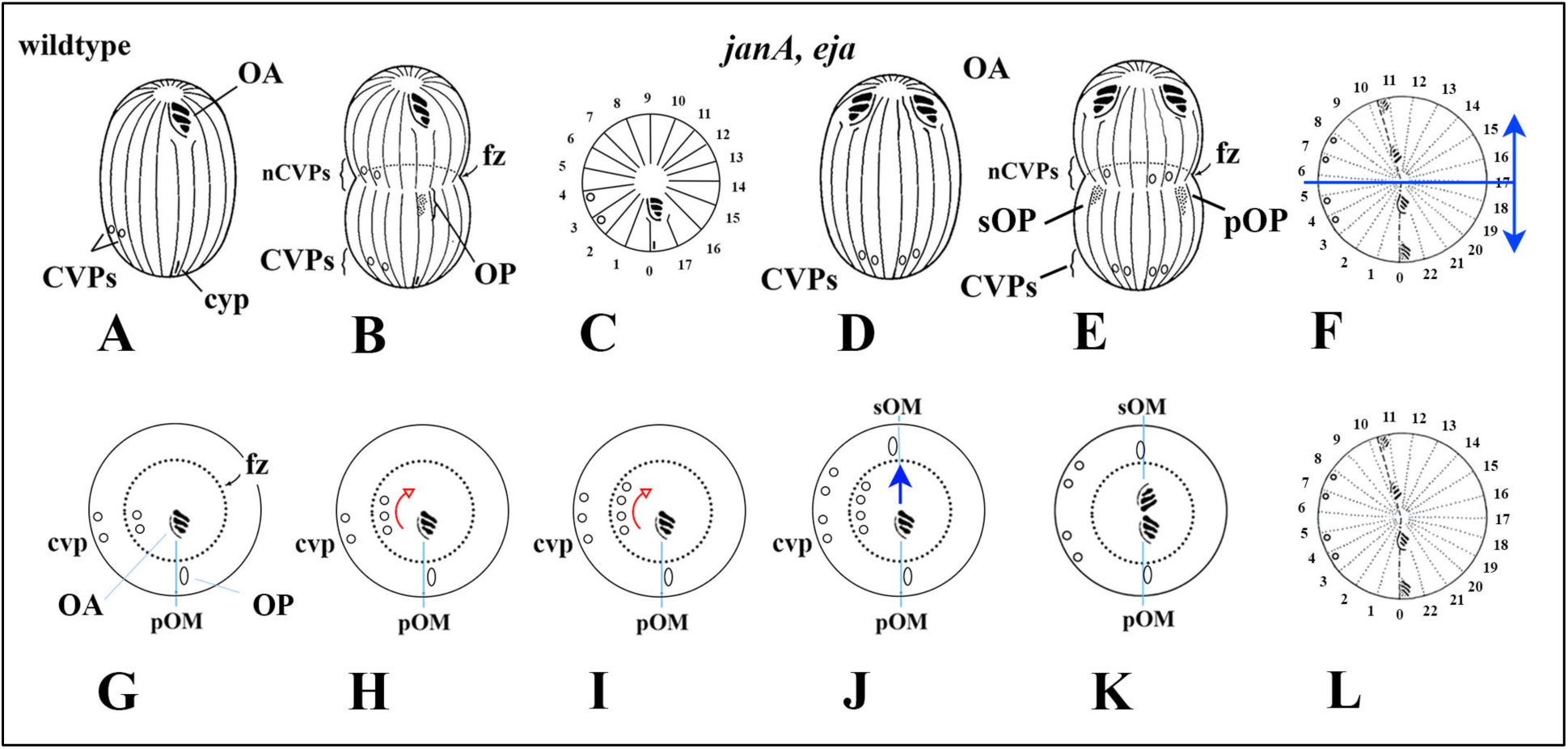
Cortical landmarks during pre-division morphogenesis in a wildtype and a *janA, eja* double-mutant. The top panel depicts: A) a ventral diagram of a non-dividing, wildtype (WT) cell; B) a ventral diagram of a WT dividing cell as new organelles are assembled, and C) a ‘polar projection’ view a non-dividing, WT cell. OA = the oral apparatus. OP = the developing (‘new’) oral primordium. FZ = the fission zone. CVPs = contractile vacuole pores. nCVPs = ‘new’ contractile vacuole pores. Linear stripes represent longitudinal rows of ciliated basal bodies. The ciliary rows are numbered in the polar view with the oral meridian (the ciliary row that gives rise to the next oral apparatus), numbered as one, and numbers increasing in a clockwise fashion looking down from the anterior pole. In the most extreme janus phenotype, the ventral pattern is duplicated with mirror-symmetry in the dorsal hemi-cell (indicated by blue arrows). The bottom panel depicts polar projections of a dividing WT cell from which the JanA gene product has been effectively removed (within an *eja*-genetic background) and the cortical phenotype observed through subsequent cell generations. The dotted circle represents the fission zone with the oral primordium assembling just posterior to it alongside the primary oral meridian (pOM). New CVPs assemble just anterior to the fission zone, and to the cell’s right of the pOM, approximately 90 degrees. The *janus* phenotype is first evident as a broadening of the CVP domain from two to three and finally four CVPs (curved red arrows). This broadening manifests in newly formed CVPs anterior to the FZ. Only after full CVP broadening does a second OP (sOP) assemble along the secondary oral meridian (sOM, blue arrow), manifesting the most extreme form of the phenotype, a mirror-image duplication of the wildtype, ventral pattern of cortical organelles.

At each cell division, one observes both the ‘inherited’ organelles of the parent cell, and newly assembled organelles. Following developmental replacement of the wildtype macronucleus with a nucleus homozygous for the *janA* mutation, cells are initially wildtype, but as cell-divisions proceed, any residual wildtype JanA gene product is diluted. Consequently, successive cell divisions reveal cortical patterning under conditions with progressively less of the wildtype product.

The result is first a broadening of the CVP domain (Figs 2 G-I) from two to three or four organelles followed by a split in the CVP domain (Fig 2 J), effectively creating two CVP domains separated by one or more ciliary rows. Finally, following the broadening and ‘duplication’ of the CVP domain, a secondary OP assembles along a secondary oral meridian (sOM) approximately 180 degrees from the primary oral meridian (pOM) (Figs 2 J,K). The fully expressed mirror-duplication only appears in 30-50% of the population, and only in combination with the 2^nd^ site enhancer mutant, *eja*.

The *janA* mutant phenotype is clearly complex, hinting at a corresponding wildtype gene with an equally complex role during vegetative development. This study identifies the *JANA* gene, documents both deletion and over-expression phenotypes, characterizes localization of the gene product, and documents a pharmacological phenocopy of the pattern mutant. Results are discussed with reference to the phenomenon of chiral, circumferential patterning in the *Tetrahymena* cell cortex (‘C-Patterning’)

## Results

### Identifying the JANA gene

*janA-1*, a recessive mutation, was mapped to the right arm of micronuclear chromosome 3 using complementation tests in crosses to nullisomic strains lacking specific micronuclear chromosomes (Bruns and Sanford, 1978; Bruns, et al., 1983; Cassidy-Hanley, et al., 1994; Frankel and Jenkins, 1979). We used ‘allelic composition contrast analysis’ (ACCA) (Jiang et al., 2017), to map the causal mutation for *janA-1*. Briefly, we sequenced pooled genomes of F2 progeny that were phenotypically either *janA-1* or wild type and the detected variants were aligned along the micronuclear reference genome. For each sequence variant, scores were computed that reflect the extent of variant linkage to *janA-1* phenotype. A peak of increased linkage was apparent on chromosome 3 around a ∼25.5 mb position (Fig 3 A, blue arrow).

**Figure 3.**
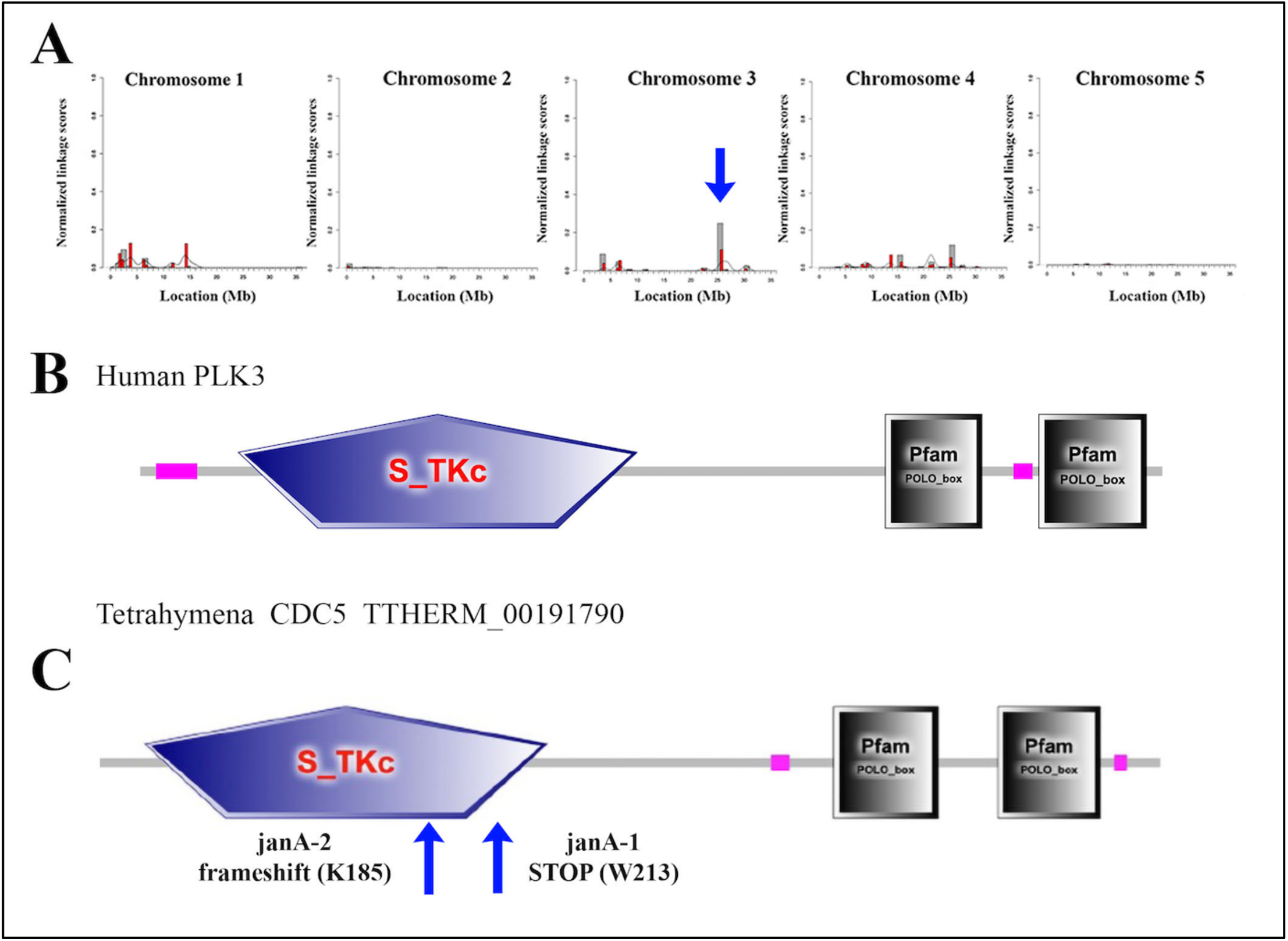
The *janA-1* alleles map to TTHERM_00191790, a gene that encodes a Polo-kinase (A) Mapping the *janA-1* mutation using the ACCA workflow of comparative whole genome sequencing. The scores reflect the degree of linkage of sequence variants to the *janA-1* phenotype among the meiotic F2 segregants. Note increased linkage signal on chr3 around the position 25.5 mb (arrow). (B,C) A comparison of the domain organization of TTHERM_00191790. Drawings generated by SMART (http://smart.embl-heidelberg.de). Arrows indicate aproximate sites of two nucleotide alterations associated with the two mutant alleles.

Within this genomic region, variant chr3:26853088 G/A was altered in 100% of reads (n=286) in the mutant pool and 99% reference in the wild-type pool (n=82). This variant affects the predicted coding sequence of *TTHERM_00191790 (also named CDC5)* by changing the W213 codon (TGG) into a TGA stop codon, resulting in the truncation of 461 amino acids (Suppl. Fig 1B). BLASTp searches revealed that the TTHERM_00191790 protein exhibits strong amino acid sequence homology to Polo kinases in other eukaryotes. A second mutation, *janA-2,* conferring essentially the same ‘*janus*’ phenotype, maps to the same chromosome arm, and fails to complement the *janA-1* mutation. Sequencing of *TTHERM_00191790* in the *janA-2* homozygote strain revealed a base pair deletion resulting in a frame shift at the K185 codon (Fig. 3, See Suppl. Fig 1B). This leads to a gain of 75 novel amino acids and premature translation termination at codon 261. The *janA-1* and *janA-2*-associated variants result in truncation of large portions of the protein including loss of a portion of the kinase domain and the POLO-box sequences, suggesting that *janA-1* and *janA-2* are both null alleles.

To further validate *TTHERM_00191790* as the *JANA* locus, we used homologous DNA recombination to delete a portion of the coding region of *TTHERM_00191790* in the macronucleus of a wildtype strain. Clones carrying a deletion within *TTHERM_00191790* were evaluated by immunofluorescence using anti-centrin and anti-fenestrin antibodies to image the cortical organelle patterns. Using anti-centrin (the BB marker), we saw no evidence of a duplicated oral apparatus or oral primordia (Fig 4A). Upon labeling with an antibody against the cortical protein fenestrin (decorating CVPs among other cortical features), we saw evidence of a weak *janA* phenotype, a broadened CVP domain, and rarely a duplicate pair of CVP domains (Figs 4 B,C). We then disrupted the *TTHERM_00191790* gene in the IA264 strain, a strain homozygous for the ‘enhancer of janA’ (*eja*) mutation. Among these cells, 30-40% of the cells exhibited the full, mirror-duplication phenotype with two OAs and OPs located 180 degrees opposite one another (Fig 4F).

**Figure 4.**
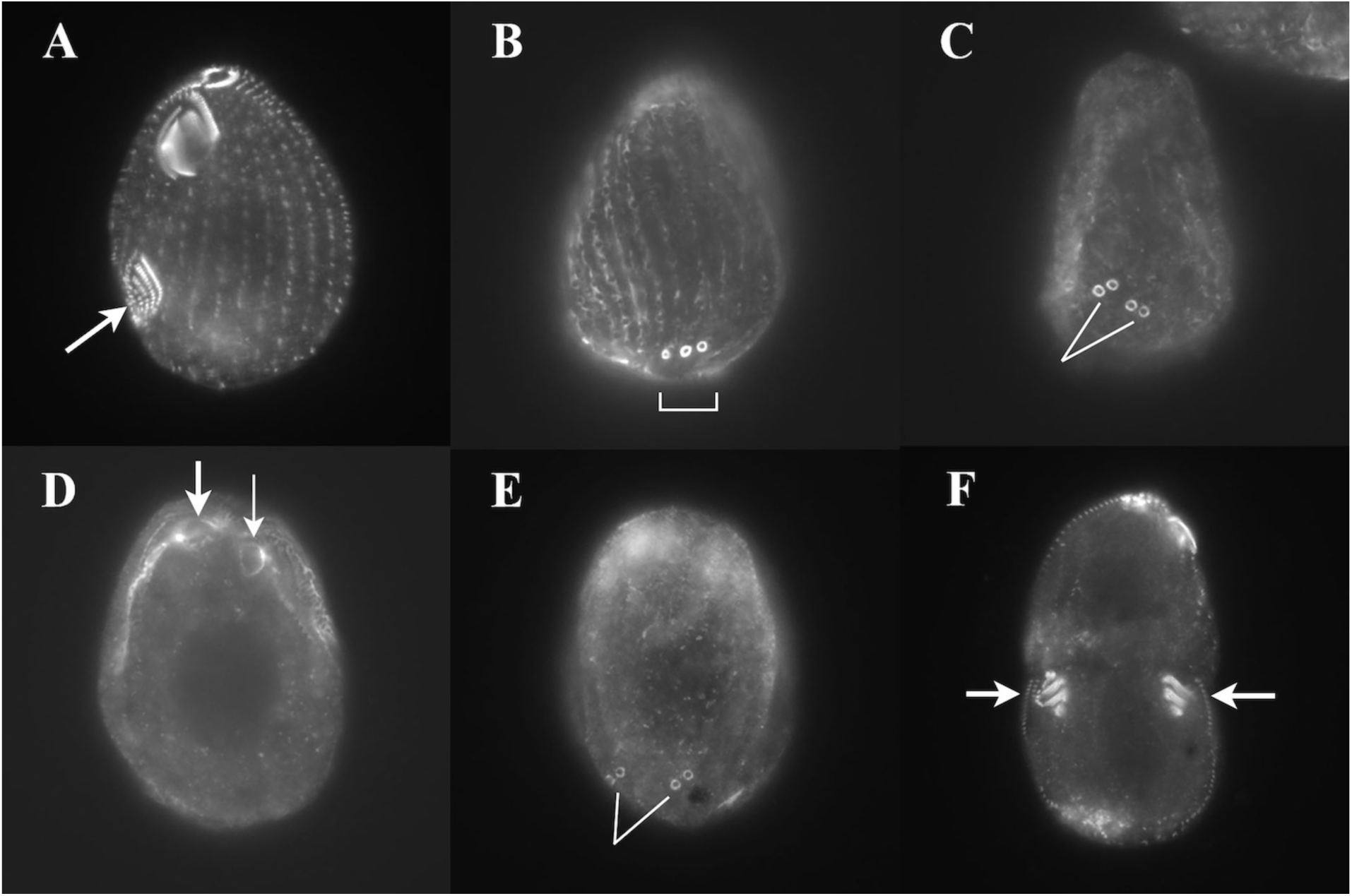
Cells in which we targeted *TTHERM_00191790* for gene-disruption and labeled with cortical antibodies. A-C). A CU428 (Wildtype) cell line labeled with A) anti-centrin antibody targeting basal bodies, we only saw cells with a single OA and OP (arrow), and B,C) an antibody to the protein fenestrin (that decorates CVPs) revealing the ‘mild’ *janA* phenotype with CVP broadening (B), bracket, and occasionally a split, dual CVP field (C), lines. When the locus was disrupted in IA264 (a cell line homozygous for *eja*) we recovered the more extreme-mirror phenotype (D-F). D, E) The same *janA* co-deletion cell labeled with antibody to fenestrin revealing twin OAs on one side (D, arrows) and twin CVP sets on the other side (E, lines). Note: fenestrin does a poor job high-lighting the OA so cells were also labeled with anti centrin revealing a proper twinned set of OAs and OPs (F arrows) representing the extreme Janus phenotype.

With these proofs: base-pair alterations in *TTHERM_00191790* gene within both *janA-1* and *janA-2*, and the full mutant phenotype exhibited by the targeted deletion of this gene in an *eja-* dependent fashion, we were confident that we had identified the *JANA* gene locus.

### JANA domain organization, phylogeny and expression profile

In addition to *TTHERM_00191790/JANA* the genome of *T. thermophila* contains 5 genes encoding closely related kinases, *TTHERM_00992960/PLK1*, *TTHERM_01443850/PLK2*, *TTHERM_01076980/PLK3*, *TTHERM_01046870/SAK1-PLK4*, and *TTHERM_00470890/PLK5* (Supplementary Fig 1A). In a phylogenetic analysis based on the kinase domains that included all 497 human sequences, *JANA*, *PLK1*, *PLK2*, and *PLK3* of *Tetrahymena* grouped in a statistically supported clade with human Polo kinases (*PLK1*, *PLK2* and *PLK3*). In *Tetrahymana*, *JANA*, *PLK1*, *PLK2* and *PLK3* all have a domain organization typical of Polo kinases, with an N-terminal kinase activity domain and a C-terminal POLO-box domain composed of two Polo Box motifs (Supplementary Fig 2). Interestingly, the coding regions of *JANA*, *PLK2* and *PLK3* have an intron at precisely the same position within the coding region (immediately downstream of codon K226 in *JANA* and at the homologous position in *PLK2* and *PLK3*) indicating that *JANA*, *PLK2* and *PLK3* are paralogs that resulted from a recent gene duplication.

In a phylogenetic analysis that included Polo kinase sequences of two additional ciliate species, *Paramecium tetraurelia* and *Stentor coeruleus*, ciliate proteins grouped in a clade containing animal *PLK1*, *PLK2* and *PLK3* and *CDC5* of *S. cerevisiae* (Supplementary Fig 3). Furthermore, *JANA*, *PLK2*, and *PLK3* form a subclade with 10 members of *P. tetraurelia* that are absent in *S. coeruleus*. *Paramecium* and *Tetrahymena* are evolutionarily more related as both belong to the class *Oligohymenophora*, while *Stentor* is a member of an early divergent class of *Heterotrichea*. Likely, *JANA*, *PLK2* and *PLK3* and the 10 *Paramecium* proteins evolved as a subtype of Polo kinases that is involved in cortical pattern formation in a subset of ciliate lineages that have an asymmetric oral apparatus located on the ventral side (see Discussion section).

Curiously, the *Tetrahymena JANA*/*CDC5* gene shows peak of mRNA abundance, not during vegetative growth and division (when the patterning occurs that led to the initial discovery of the mutant), but during conjugal development (Supplementary Fig. 4A). In particular, there is a peak of expression during meiosis (2-4 hrs into mating), and during macronuclear anlagen formation (between 6-12 hours). During vegetative growth, the *JANA*/*CDC5* gene shows elevated expression just before cytokinesis (Supplementary Fig 4B).

### JanA marks the left-dorsal region of Tetrahymena cortex

The *JANA* gene was modified to encode a C-terminal GFP fusion protein by editing the endogenous *JANA* locus. In vegetatively-growing cells, JanA-GFP was associated with basal bodies exhibiting a dorsal-ventral gradient of fluorescence intensity, with highest concentration in the left-dorsal hemi-cell from the 6^th^ ciliary row, to the 15^th^ ciliary row (Fig 5E). Ventral basal bodies were also labeled, though much less intensely. In addition, JanA was detected in the contractile vacuole pores (arrows, Fig 5C). It is notable that the cortical domain most richly occupied by JanA:GFP overlaps the region where the mirror-duplication of ventral organelles is manifest in the loss-of-function *janA-1* mutants.

**Figure 5.**
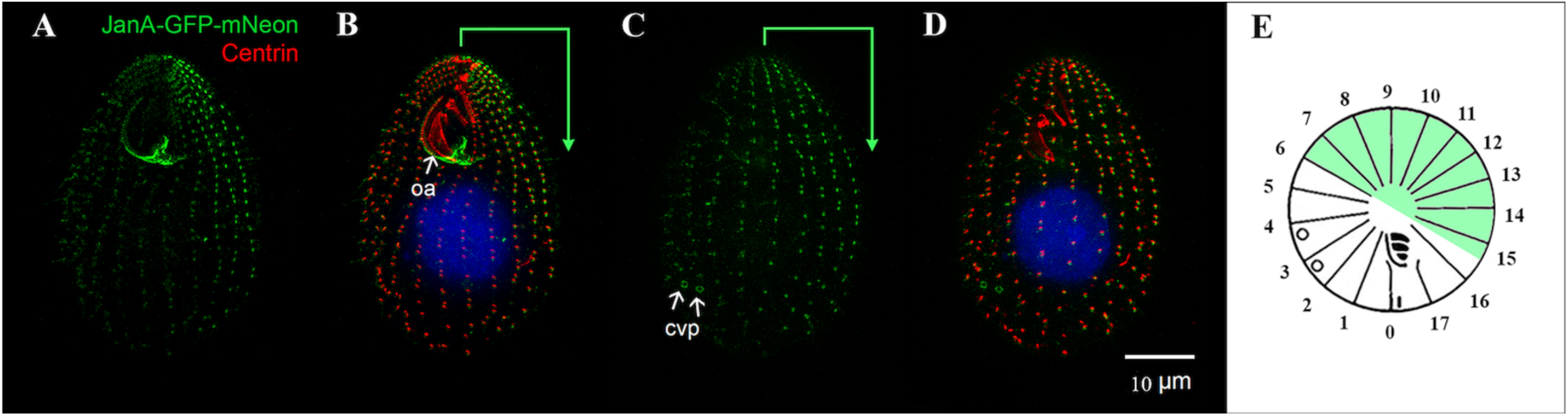
*Tetrahymena* cells expressing a JanA:GFP fusion protein from the endogenous promoter. Cells were fixed, and GFP-localization was enhanced using an antibody to GFP with a FITC-tagged secondary antibody (green). Cells were counter-stained using antibody to centrin with a Cy3 secondary antibody (red). A) A ventral view of a JanA:GFP labeled cell. B) The same cell showing anti-centrin (red) and JanA:GFP enriched around basal bodies in the ‘left-dorsal hemi-cell’ (bracket). C, D) The same cell viewed from the dorsal side. Arrows highlight CVPs decorated by JanA:GFP. E) Polar projection showing the left-dorsal hemi-cell (green) within which JanA:GFP protein appears more concentrated, and CVPs on ciliary rows 3 and 4.

### Other PLK:GFP gene products mark distinctive, alternative cortical domains

GFP-tagging of PLK1, PLK2, PLK3, and PLK5 revealed distinctive cortical localizations centered around various fields of basal bodies (Fig 6). PLK1 shows uniform circumferential distribution with a conspicuous anterior-posterior gradient, especially visible during cell division just posterior to the fission zone. PLK2 exhibits a pattern of localization that mirrors that of JanA. PLK2 is high over basal bodies on the right ventral hemi-cell, and impoverished in the left dorsal hemi-cell where JanA is abundant. PLK3 decorates all the cell’s basal bodies with enhanced brightness around the ventral ciliary rows that give rise to CVPs (arrows, Fig 6K). PLK5 decorates all the basal bodies, with possible intensification in the ventral hemi-cell. These observations suggest that most if not all of *Tetrahymena* PLks may be involved in cortical patterning acting in specific subregions of the cell circumference.

**Figure 6.**
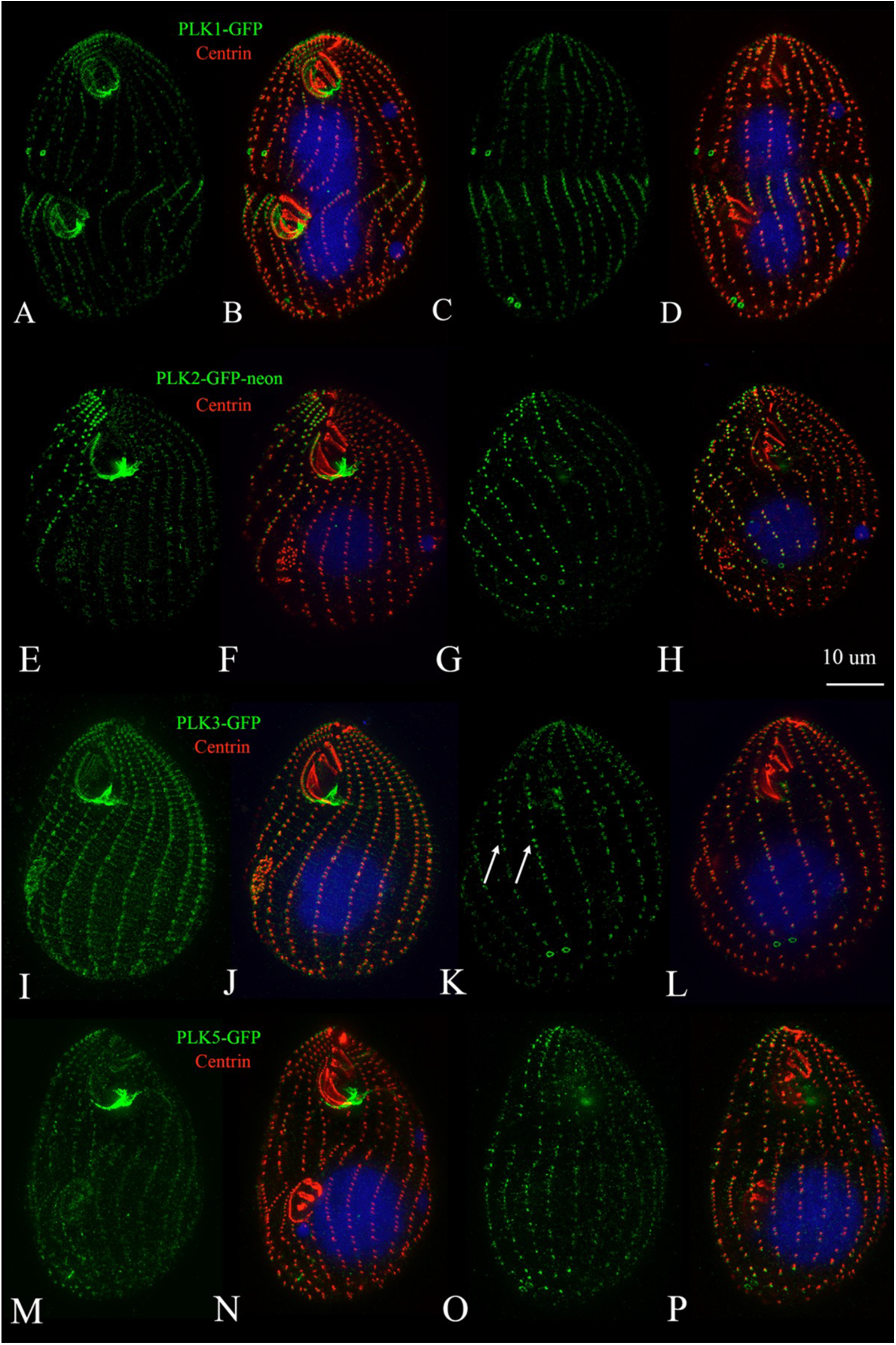
PLK localizations for *Tetrahymena* PLK1,2,3,5. PLK genes were expressed from endogenous promoters, and tagged with either GFP or GFP-neon. GFP signals were enhanced with polyclonal antiserum to GFP. Basal bodies were counter-stained with an antibody to centrin (red).

### Prolonged over-expression of GFP-JanA severely disturbs the cortical pattern

We edited the *JANA* gene to place the coding sequence under control of the cadmium-inducible *MTT1* promoter (Shang *et al*., 2002), and analyzed the consequences of GFP-JanA overproduction during vegetative propagation (Fig 7). The *MTT1* promoter is ‘leaky’ when cells are raised in iron-rich Neff medium, driving a modest level of expression even without addition of cadmium ions. With no cadmium added, we observed the same dorsal pattern of basal body localization observed using the endogenous promoter (as seen in Fig 5A). When cells carrying the MTT1-GFP-*JANA* transgene (with an N-terminal tag) were exposed to cadmium chloride (2.5 μg/ml) for 2-4 hrs, fluorescence intensified over both dorsal and ventral basal bodies, the developing oral primordium (OP), the mature OA, and the CVPs. All these structures contain microtubules and therefore it is likely that JanA either binds directly or is in a complex with another protein that has a microtubule-binding domain. This inducible, GFP-*JANA* expression cassette offers a potentially powerful tool for future studies of cortical architecture, pattern formation and development in live *Tetrahymena*. Cells exposed to cadmium for 2-4 hrs appeared phenotypically normal (Fig 7). Prolonged expression of the GFP-fusion gene (beyond 6 hrs) led to a progressively more severe over-expression phenotype. After two hours of induced expression, cells undergo ‘***oral replacement***’ (Fig 8A), an alternative developmental pathway during which a new OA forms in the proximity of the old OA.

**Figure 7.**
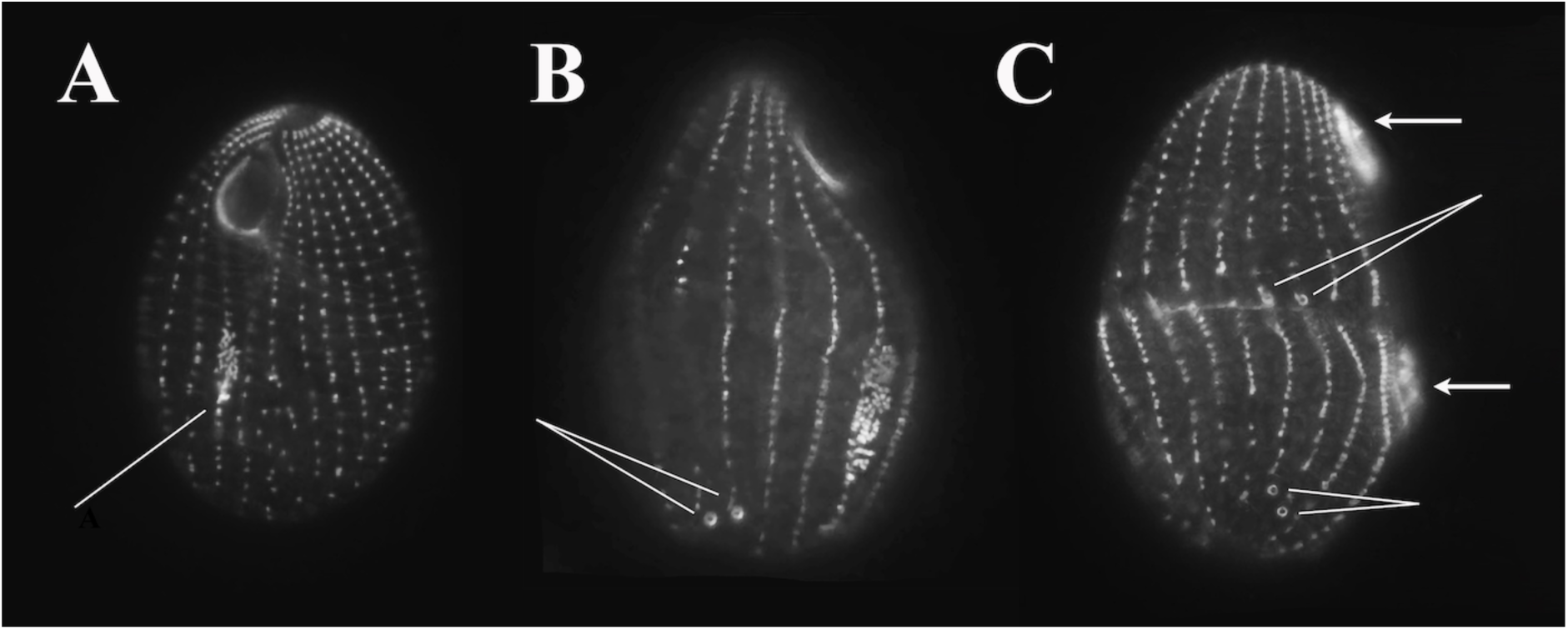
Wildtype (CU428) cells expressing the inducible GFP-JanA fusion protein following 4 hours of exposure to 2.5 ug/mL cadmium chloride. A) All the basal bodies (dorsal and ventral) are now decorated including the basal body proliferation field that marks assembly of the OP (thin arrow). B) A similarly labeled cell undergoing pre-division oral morphogenesis, showing the typical pair of CVPs (thin arrows). C) A slightly later stage in pre-division morphogenesis showing the fully developed OP and OA (thick arrows) and the old and new CVPs (thin arrows), new CVPs forming just anterior of the developing cytokinetic furrow.

**Figure 8.**
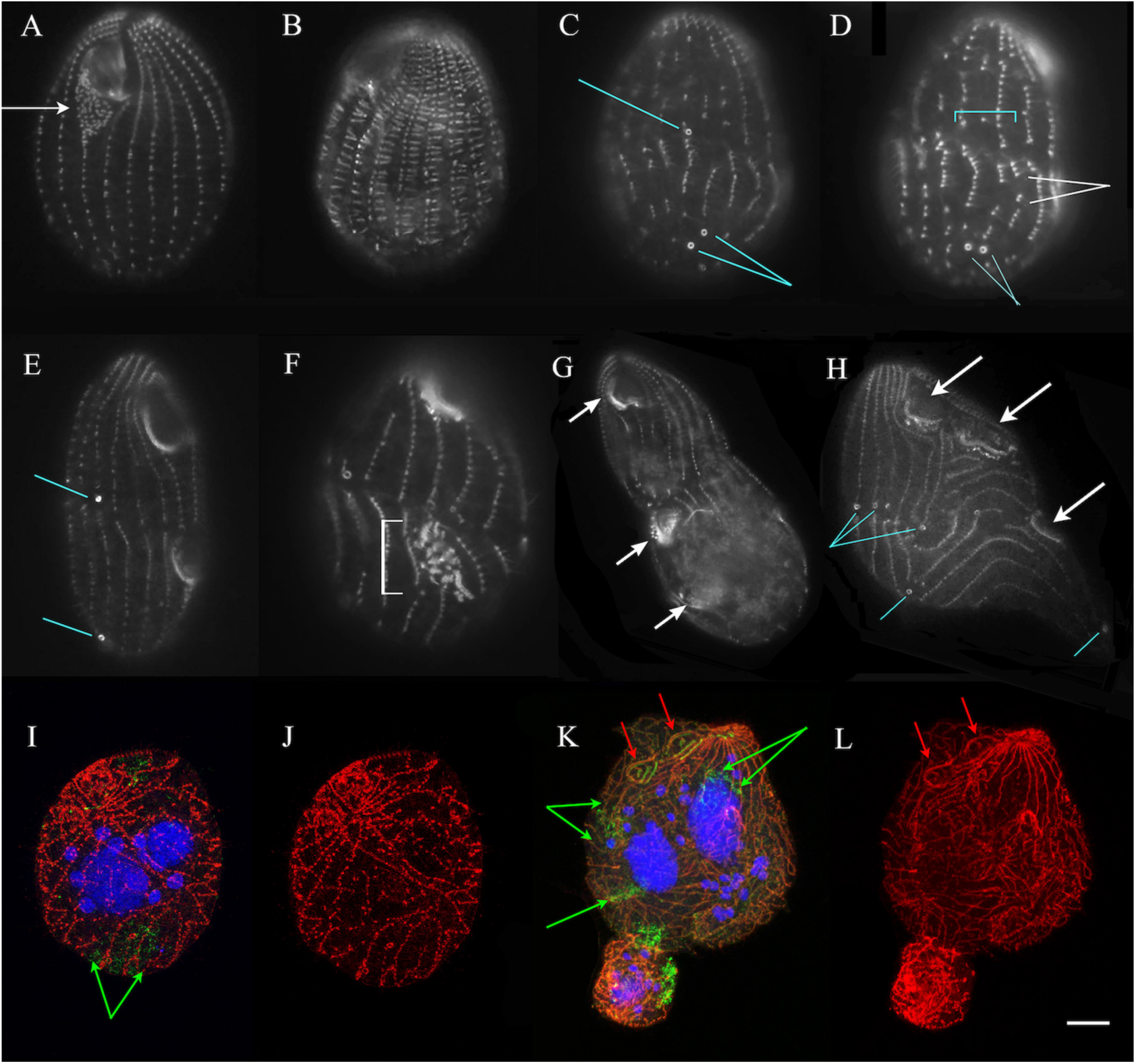
Wildtype (CU428) cells expressing the inducible GFP-JanA fusion protein following 5-28 hours of exposure to 2.5 ug/mL cadmium chloride. A) After 5 hrs, we witnessed a wave of ‘oral replacement’, a process typically triggered by starvation. B) After 6-8 hrs. of cadmium-induction, GFP-janA decorates the transverse microtubules. C, D) Cells that divided after 6-8 hrs of induction began to exhibit a reduction in the breadth of the CVP domain (turquoise lines) from the typical two to one (Fig. C) to zero (Fig. D, bracket). Breaks also began to appear in the ciliary rows of basal bodies (white pointers). E) Fenestrin-labeled cells incubated overnight with cadmium exhibited a reduction in both the mature CVP domain (lower pointer) and the developing domain (upper pointer). F) Centrin-labeled cells began to exhibit disruptions in oral assembly (bracket). Membranelles appear to be forming in a variety of orthogonal relationships with no distinct UM. G, H). Overnight induction of the GFP-JanA fusion protein produced cells arrested in cell division. G) A centrin-labeled cell showing three OAs (arrows) and one failed fission zone. H) A cell with 3 OAs (white arrows) and 3 CVP fields (turquoise lines). I-L) *Tetrahymena* cells following 8-24 hours of induced over-expression of the GFP-JanA fusion protein. Red indicates the GFP-JanA fusion protein highlighted with an antibody to GFP. Green = antibody to centrin decorating basal bodies and the contractile vacuole ‘complex’. Blue = DAPI staining of supernumerary MICs and MACs. K and L depict a ‘terminal phenotype’ in which the MAC has divided once, two OA are conspicuous (red arrows) and there are numerous MICs. (Scale bar = 10 um).

Oral replacement (OR) is typically seen in wild-type cells that have been subjected to nutrient deprivation (Frankel, 1969; Nelsen, 1978; Williams & Frankel, 1973). OR is initiated as the posterior half of the undulating membrane (UM) and the anterior-most basal bodies of the right post-oral ciliary row become disorganized, forming a replicative field of basal bodies that eventually establishes a new replacement OA. The population of cells that undergo OR in response to GFP-JanA over-expression was quite high, 50% after 5hrs. This suggests, that virtually the entire population carries out OR in response to GFP-JanA over-expression over the course of 4-6 hrs.

After four hours of continuous over-expression, we began to observe a suite of more severe abnormalities (Fig 8). First, GFP-JanA localization spreads from the basal bodies to the transverse microtubules (Fig 8 B). At the same time, cells undergoing cell-division begin to exhibit a reduction in the breadth of the new CVP domain from 2 to one, and even zero, though such examples were only observed in the anterior division product prior to cytokinesis (Figs 8 C-E). Daughter cells that complete cell division lacking a CVP are presumably unable to osmoregulate and die. Cells began to exhibit fragmentation of the ciliary rows (Fig 8 D) and with prolonged GFP-JanA over-expression, the OP exhibits signs of pattern derangement (Fig 8 F). After six hours, over-expression cells exhibit difficulties with cell division, especially MAC division and cytokinesis. Curiously, MIC division and stomatogenesis seem relatively unaffected resulting in cells with multiple OAs, multiple MICs, but only one or at most two MACs (Figs 8 G-H). Between 8 and 24 hrs of cadmium-induced over-expression, and following multiple failed rounds of cytokinesis, some cells assume a configuration in which there appears to be one anterior pole and multiple posterior poles (Fig 8H). Though it is attractive to postulate that over-expression of GFP-JanA affects mechanisms that regulate anterior/posterior polarity in the cell, we suspect this result is more likely an attempt on the part of the cell, to re-integrate multiple (failed) division products such as those seen in Fig 8G. Continued GFP-JanA over-expression results in some rather spectacular cyto-geometries, sometimes with ciliary rows folding back on themselves highlighting the continued replication of basal bodies despite fission failure.

Under continuous exposure to cadmium-driven GFP-JanA over-expression (24 hours or longer), cells undergo cell division cycles, while exhibiting cytokinesis failure. Typically, these ‘monster’ phenotypes exhibit two MACs and two OAs, with a multitude of MICs and an excess of basal bodies (Fig 8I-L). Curiously, one of the final consequences of the over-expression phenotype, appears to be the loss of centrin from the somatic basal bodies. Antibodies to centrin normally decorate the basal bodies that support the somatic and oral cilia, as well as a diffuse cytoplasmic cloud associated with the contractile vacuole system. Using the green CV assemblage as a positive control, as well as the persistent green centrin label over the mature oral membranelles, we confirmed that, though the GFP-JanA marker remains in linear punctae associated with ciliary rows, these punctae are now negative for centrin labeling.

### Volasertib, a PLK inhibitor, broadens the CVP domain

Having identified *JANA* as a Polo kinase, a Ser-Thr kinase homologous to the cell-cycle regulator in vertebrates, we were curious to learn if JanA was sensitive to drugs developed as potential anti-cancer treatments that antagonize PLK activity (Steegmaier, et al., 2007; Gjertsen, & Schöffski 2015). We tested two of these: volasertib and BI2536. Cells were cultured overnight in a range of drug concentrations, and cortical features were examined following brief expression of the MTT-driven *JANA*:*GFP* fusion gene. CVP numbers provide a convenient measure of the strength of the drug effect, and oral organization allows assessment of the more extreme phenocopies. Figure 9 shows the dose-response for varying concentrations of volasertib on cortical patterning as assessed by CVP count.

**Figure 9.**
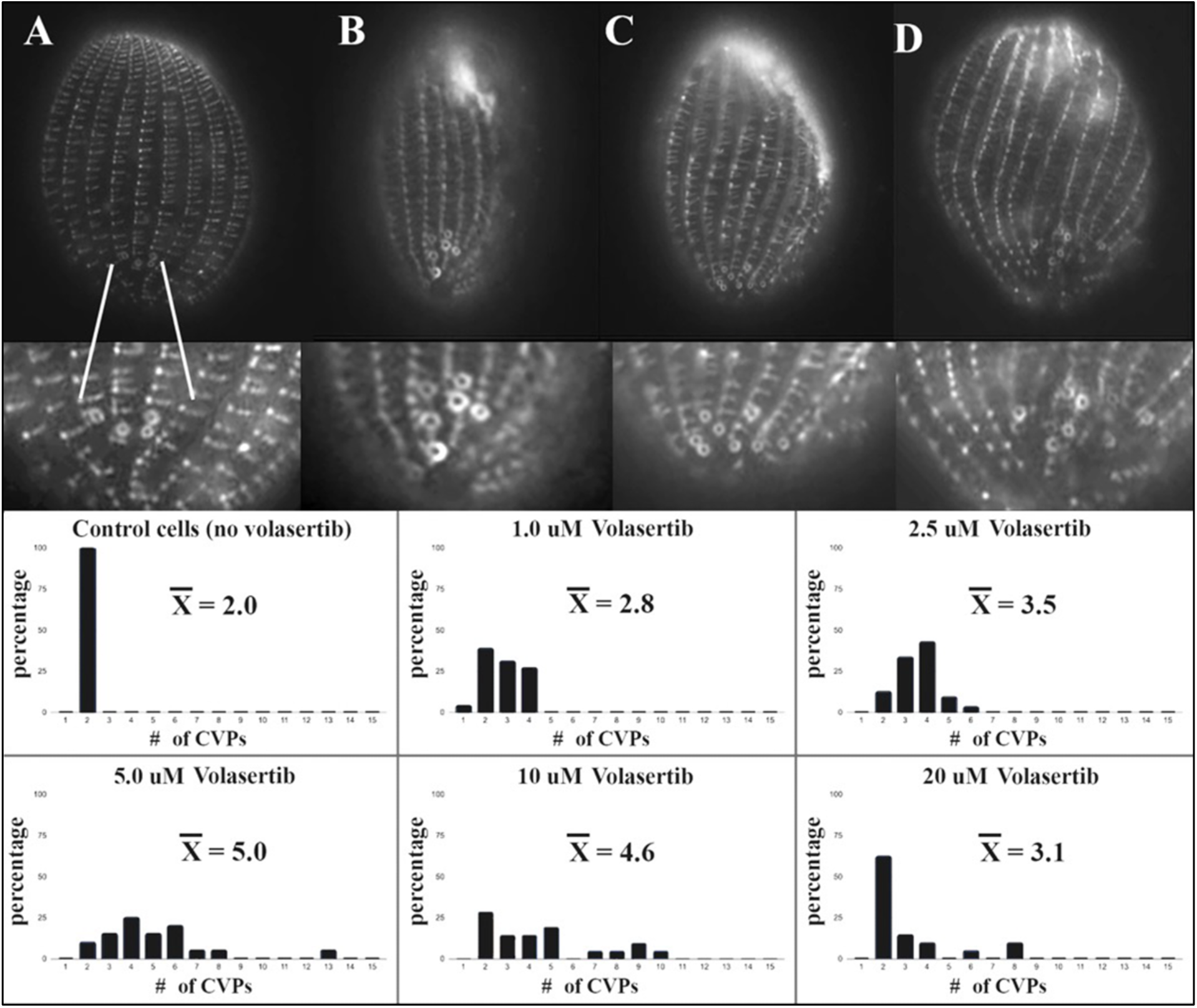
Wildtype cells (CU428) treated overnight to a range of volasertib concentrations, and imaged following brief expression of the MTT:GFP-*JANA* fusion gene with 2.0 ug/mL CdCl_2_. Four examples from the 5.0 uM treatment are shown expressing four, six, eight and ten CVPs (Figs. A-D). 5.0 uM volasertib gave the most consistently high number of CVPs with one example expressing 13 CVPs instead of the typical two.

The CVP phenotype expressed in volasertib-treated cells was not what we expected. When the JanA gene product is progressively diminished by developmentally-induced expression of the mutant allele in the wild-type background, we see a modest broadening of the CVP domain from 2 CVPs to 3 or occasionally 4. This broadened field of CVPs often becomes divided by a gap of one unoccupied ciliary row (Fig 2). Volasertib creates a profound expansion of the CVP-domain, more than doubling its average width at the optimal concentration (5.0 μM), and producing examples with as many as 13 CVPs, a phenotype never observed in *janA* mutants. Curiously, the CVP-broadening phenocopy became less severe at higher concentrations. We suspect that this is due to progressively more severe suppression of cell division at higher concentrations, preventing assembly of new CVP fields under the influence of the drug. In such cases one sees, increasingly, the persistent ‘parental’ CVP domain of non-dividing cells.

Only rarely did drug-treated cells exhibit a second OP developing 180 degrees around the dorsal surface in response to volasertib. It is noteworthy, that the volasertib-treatment didn’t simply ‘broaden’ the cell’s CVP domain (causing CVPs to assemble alongside an increased number of ciliary rows), the CVP domain was expanded along the anterior-posterior axis of a ciliary row as well, with 2-3 CVPs seen alongside a single kinety (Fig 9 B). Hence, it is more accurate to state that volasertib <u>expands</u> the CVP domain both laterally (left-right) and longitudinally (anterior-posterior). These results suggest that the *janA* mutation is partially, (though imperfectly) phenocopied by inhibiting all PLK activity. That said, broad-spectrum inhibition implicates general PLK activity in the control of the dimensions of the CVP domain, one component of the *janA* phenotype.

### Pharmacological effects of BI2536, another PLK inhibitor

BI2536 produced an even stronger expansion of the CVP domain, with occasional cells exhibiting the janus global mirror-pattern duplication (Figs 10, 11). The mirrored cortical pattern shown in Fig 11 A-D, is remarkable in its similarity to the *janA* mutant phenotype suggesting that BI2536 is a more potent PLK inhibitor than volasertib in *Tetrahymena*, and is capable of producing a full janus phenocopy. Note: the images in Fig 11 A and B are through-focus images of <u>the same cell</u> revealing both a dorsally and ventrally located OA. Again the drug-induced phenocopy is unlike the *janA* phenotype in the sheer over-abundance of CVPs (Figs 10, 11 F) and in the unexpected multiplication of the cell’s cytoprocts. Wildtype cells have a single cytoproct (Fig 11 E), whereas BI2536-treated cells display 4 or 5 (Fig 11 F).

**Figure 10.**
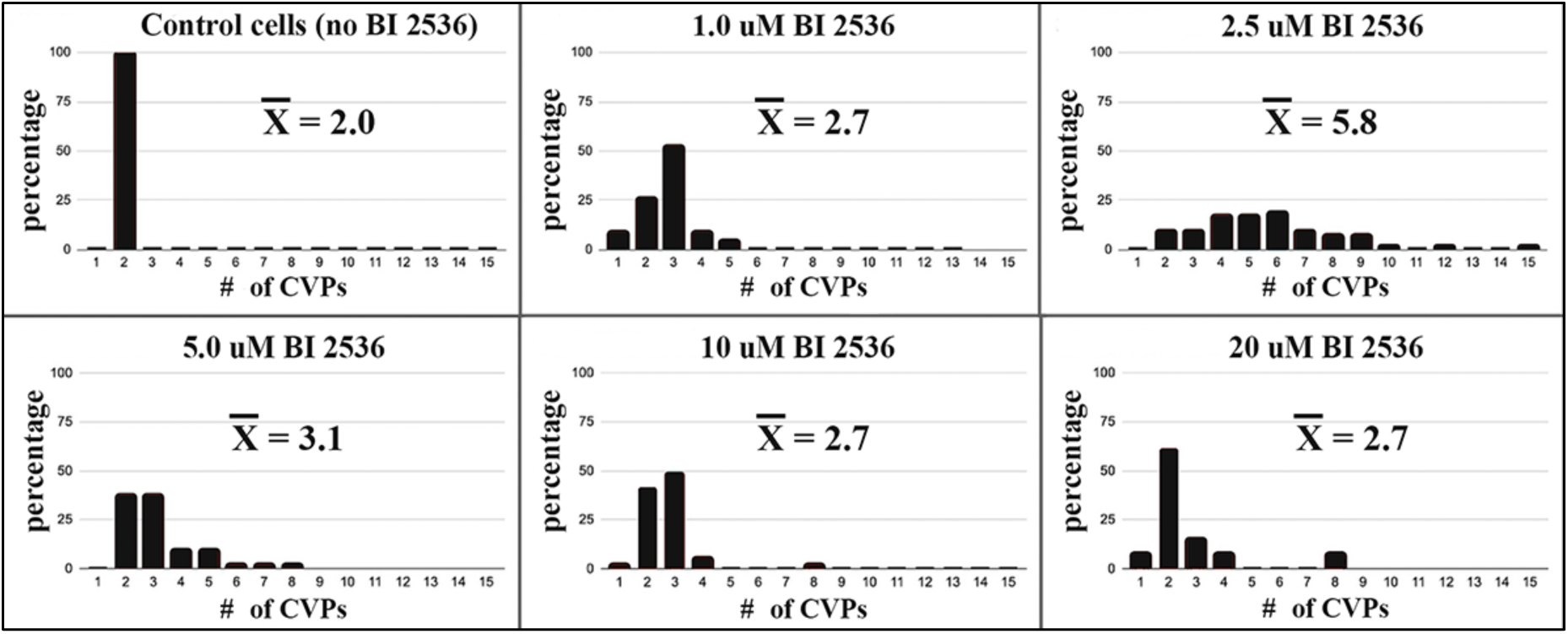
Wildtype cells (CU428) treated overnight to a range of BI2536 concentrations, and imaged following brief expression of the MTT:GFP-*JANA* fusion gene with 2.0 ug/mL CdCl_2_. 2.5 uM BI2536 gave the most consistently high number of CVPs with one example expressing 15 CVPs instead of the typical two.

**Figure 11.**
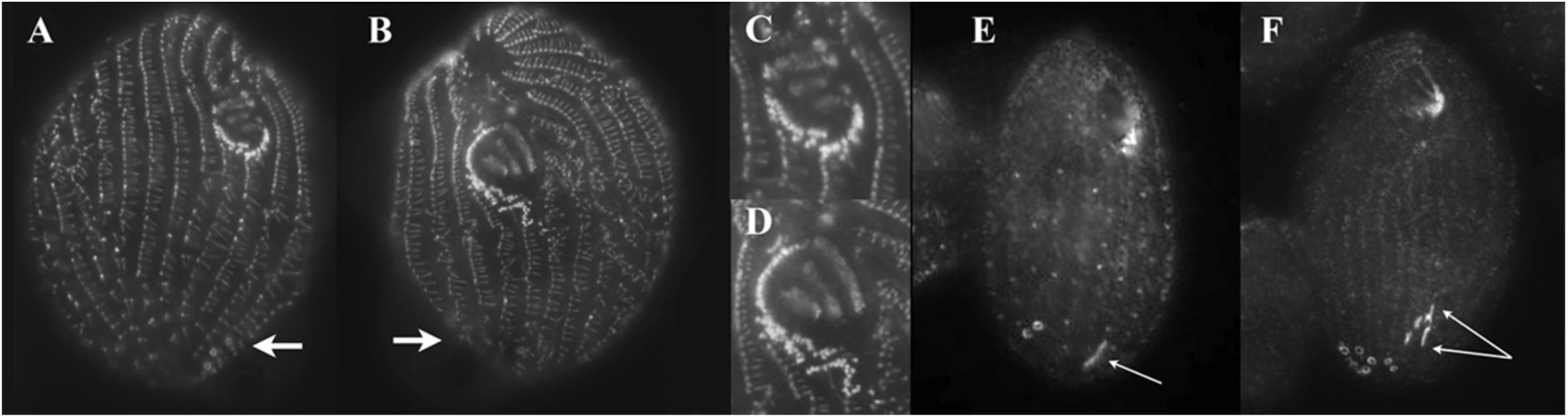
*eja* cells expressing the GFP-JanA fusion-protein and grown under the influence of 2.5 mM BI2536. A) The dorsal view, showing an OA with partial reversal of local asymmetry (expanded view in C). A CVP domain is visible lower right (arrow). Notably, this CVP domain is on the cell’s left of the secondary OA and to the cell’s right of the primary OA. B) The ventral view showing a more normal primary OA. Both OAs exhibit some evidence of UM disassembly typical of oral replacement. D) An expanded view of the primary OA seen in Fig B. Note, we used the intrinsic asymmetry of the transverse microtubules for orientation, printing images as they would appear from each surface, rather than the reverse image one would observe of the back of the cell as one down-focuses through the specimen. (Transverse microtubules extend to the cell’s-left, viewer’s right of each ciliary row). E) A wildtype cell labeled with antibody to fenestrin (a protein that decorates CVPs and the Cytoproct: Cyp). Arrow indicates the single Cyp. F) *eja* cells expressing the GFP-JanA fusion-protein and grown under the influence of 2.5uM BI2536. This cell exhibits 7 CVPs and at least four Cyps (arrows).

### PLK inhibition by BI2536 creates a more extreme phenocopy in cells homozygous for eja

The *janA* phenotype is enhanced when expressed in cells homozygous for a 2^nd^-site phenotypic enhancer, *eja*. We compared the effect of 2.5 μM BI2536 on CU428 cells (without the *eja* mutation) with that on IA264 cells (cells homozygous for *eja*) (Fig 12). BI2536 produced almost 1.5 times more CVPs in cells that were homozygous for the *enhancer of janA* mutation.

**Figure 12.**
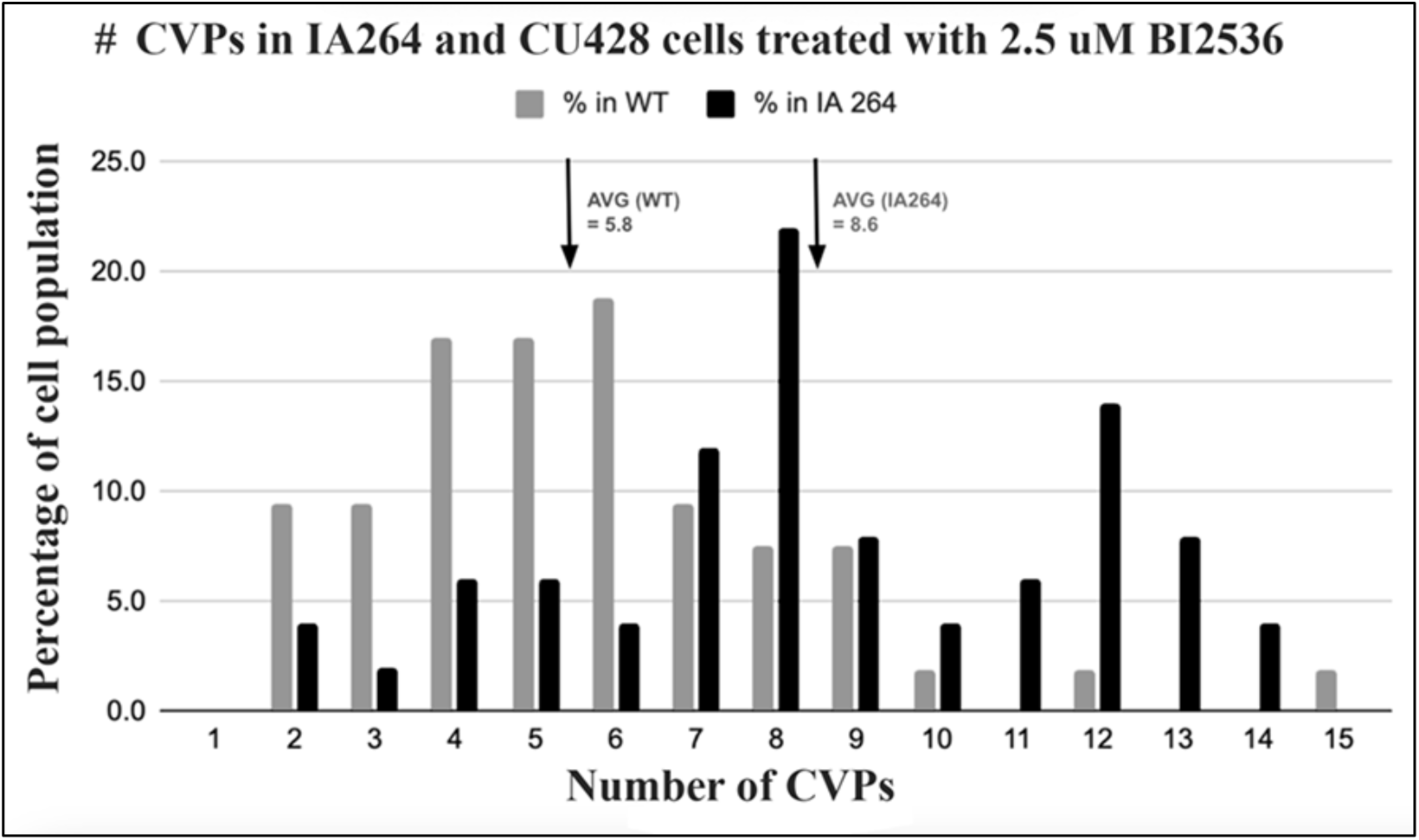
CVPs were counted in two different cell lines treated overnight with 2.5uM BI2536. CU428 was used as a wildtype control and the IA264 cell line is homozygous for the *eja* mutation. BI2536 had a significantly greater effect on CVP domain-broadening in the *eja* mutant (T-test result: p< 1.3 e^-6^).

We also observed abnormal features in the OAs of BI2536-treated *eja* cells (Fig 13). OAs displayed a disorganized field of basal bodies where the tight, double rowed undulating membrane (UM) should be, and frequently adopted a local-mirrored configuration of primary membranelles 1-3 (Figs 13 C-E). This same mirrored configuration was seen in BI2536-treated wildtype cells, though to a lesser extent. The mirrored-membranelle form, though highly consistent in treated *eja* cells, is difficult to interpret. In such cells, the UM appears to have dissociated into an anarchic field of basal bodies (Fig 13 B) reminiscent of the oral replacement primordium (ORP) described earlier following JanA-over-expression (Fig 8A).

**Figure 13.**
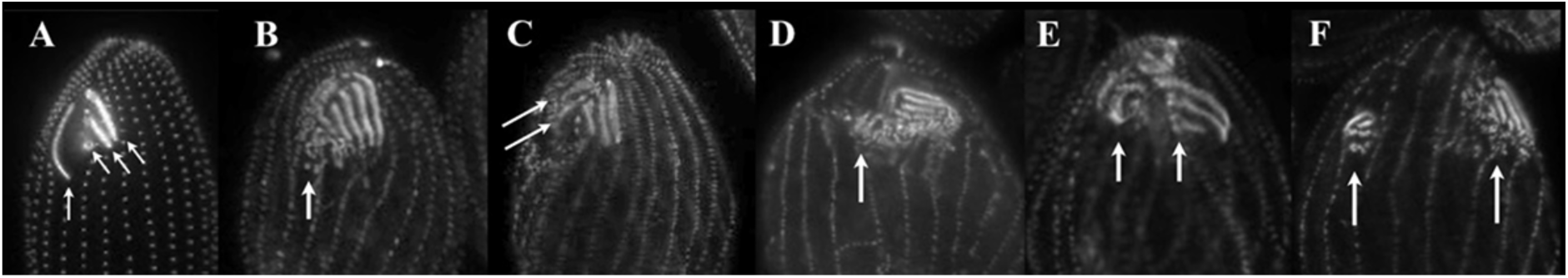
The effect of BI2536 on the OA of IA264 cells (homozygous for *eja*). A) A wildtype OA labeled with Poc1:GFP (photo courtesy of Chad Pearson). Vertical arrow is pointing to the UM (undulating membrane). Diagonal arrows indicate membranelles 1,2 and 3. B) IA264 cell exposed to BI2536 showing a dispersed field of basal bodies where the UM should be. C) Basal bodies in the region normally occupied by the UM appear to be organizing into rows angling upward from lower left to upper right (arrows). D) Appearance of what appears to be a 2^nd^ OA (arrow) to the viewer’s left of the primary OA, with membranelles angling from lower left to upper right. E) Even more discrete secondary OA (left arrow). Note, primary OA (right arrow) is still lacking a UM. F) Two discrete OAs separated by three ciliary rows. The primary, right-most OA still lacks a well-organized UM.

During OR, the UM dissociates, creating a field of basal bodies. These basal bodies join others proliferating from the anterior-most region of the first post-oral ciliary row. In wildtype cells, this proliferating ORP field of basal bodies (similar to the OP that forms at midbody during pre-division morphogenesis), organizes into a new OA located posterior to the original OA. The latter undergoes disassembly and presumably, endocytic resorption.

In BI2536-treated *eja* cells, this loosely organized field of basal bodies frequently assembled into what appears to be a 2^nd^ OA situated lateral to the primary, newly assembled OA, (Figs 13 C-E) and assembled with superficial mirror-symmetry to the first. Membranelles of the second OA often angle from the cell’s lower right to upper left, rather than normally from upper right, to lower left. UM disruption coupled with a mirrored-symmetry of the membranelles was observed for both dorsal and ventral OAs in the full-janus phenocopies as well (Figs 11 A-D).

## Discussion

### A model explaining the role of the JanA/CDC5/PLK protein in global, cortical patterning

Three observations from this study provide insight into the mechanism of circumferential patterning (C-patterning) within the *Tetrahymena* cell cortex. Loss of *JANA* gene function produces a global, mirror-duplication of organelles normally assembled in the ventral hemi-cell within the dorsal cell cortex. Transformation of the dorsal hemi-cell (devoid of the major cortical organelles) into a mirrored pattern of the ventral organelles, represents a novel example of intracellular, cortical homeosis. Endogenously driven JanA-GFP is concentrated around basal bodies of the left-dorsal hemi-cell. Over-expression of GFP-JanA diminishes (or eliminates) the CVP domain, and can disrupt OA development as well. It is tempting to invoke a role for the wildtype JanA gene product in suppressing a cryptic mirror-pattern of the ventral organelles within the dorsal cortex. This would suggest that the mirrored-pattern of cortical organelles represents a ‘ground state’ or default option that is normally suppressed in the wildtype cell.

### A model for regulating the pattern of organelle assemblages in the Tetrahymena cortex

A working model appears in Fig 14. We cautiously chose to model CVP and OP patterning using separate morphogens. In this model, we invoke a ventral morphogen (depicted in pink) centered at 9:00 and reaching from 6:00 to 12:00 viewing the polar projection as a clock-face. This right-ventral Oral Morphogen drives OP assembly at its margins possibly requiring interaction with a complementary left-dorsal morphogen, a situation that only occurs at their margins or interface. JanA (green) overlaps the Oral Morphogen between 9:00-12:00, and serves to repress organelle assembly (Fig 14A). Loss of JanA due to mutation, allows OP-induction at this normally cryptic site (Fig 14B). These are all observed phenomena. For CVP patterning, we propose a similar cortical morphogen (turquoise in Figs 14C-E). This morphogen, also centered at ∼9:00, licenses CVP assembly along neighboring ciliary rows. Again, the dorsal, JanA morphogen (green) overlaps this unidentified CVP morphogen at its right margin (∼10:00), and serves to repress organelle assembly (Fig 14C). Loss of JanA due to mutation, allows the CVP-induction to spread (Fig 14D), whereas JanA over-expression (Fig 14E) represses CVP induction even within the normal cortical locations (Refer to Figs 8C,D). These are also observed phenomena. This model predicts the existence of left-ventral morphogens that drive CVP and OP assembly.

**Figure 14.**
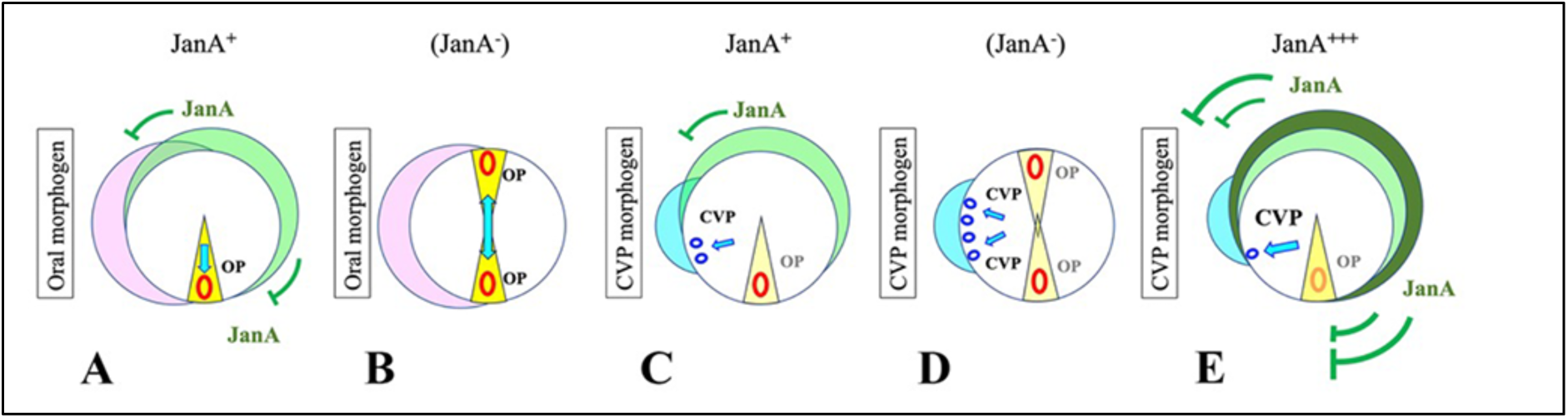
A model for how JanA controls circumferential patterning in *Tetrahymena*. A-B) The role of JanA in determining OP assembly sites. A) We hypothesize a right-ventral morphogen (pink) determines the location of oral assembly at its margins (yellow wedge/ red oval). JanA (green) represses Oral assembly in the dorsal location (12:00), and permits oral assembly at the ventral location (6:00). B) In the absence of JanA repression, oral morphogenesis is initiated at both dorsal and ventral locations. C-E) The role of JanA in determining CVP assembly sites. We hypothesize a separate, CVP-inducer (turquoise). C) JanA (green) is hypothesized to antagonize CVP-induction in the areas of turquoise/ green overlap. D) In the *janA* mutant, the turquoise CVP inducer exhibits a broader area of cortical influence, expanding the CVP domain. E) Cells over-expressing JanA, exhibit reduced CVP assembly (and possible, modest disruption of oral assembly).

### Modeling the impact of broad-spectrum PLK inhibitors on cortical development

*Tetrahymena* have 6 Polo kinases including JanA. The PLK inhibitors we deployed (volasertib and BI2536) are notable for their general efficacy in suppressing PLK activity and likely interfere with multiple PLK pathways within our model organism (Steegmaier, et al., 2007; Gjertsen, & Schöffski 2015). Consequently, drug effects on cortical morphogenesis cannot be specifically assigned to repression of the JanA kinase alone. Several features of the phenocopy produced using these drugs are notably different from the *janA* loss-of-function phenotypes. First, PLK inhibition produces a profoundly expanded field of CVPs. In the *janA* mutants (both point mutations and the knockout strains), three or at most four CVPs assemble in side-by-side arrangement, or split into two fields separated by a ciliary row. Cells exposed to PLK inhibitors express up to 15 CVPs, often assembling in vertical (anterior-posterior) clusters, two to three deep along a single ciliary row. These observations suggest that multiple drug-sensitive PLKs are involved in patterning the CVP domain, exerting an inhibitory influence over the dimensions of the organelle assembly platform. We also observed multiplication of the cytoproct (Cyp) in cells under PLK inhibition.

There was also a curious derangement of the primary oral apparatus in cells treated with PLK inhibitors not seen in the *janA* mutants. The UM appeared in a state of disassembly with basal bodies only loosely associated. These loose basal bodies frequently assembled into a second set of oral membranelles that were organized with mirror-symmetric orientation to the first: a ‘local’ mirrored-symmetry affecting orientation within this complex cortical organelle. A model depicting local membranelle symmetry derangement appears in Fig 15.

**Figure 15.**
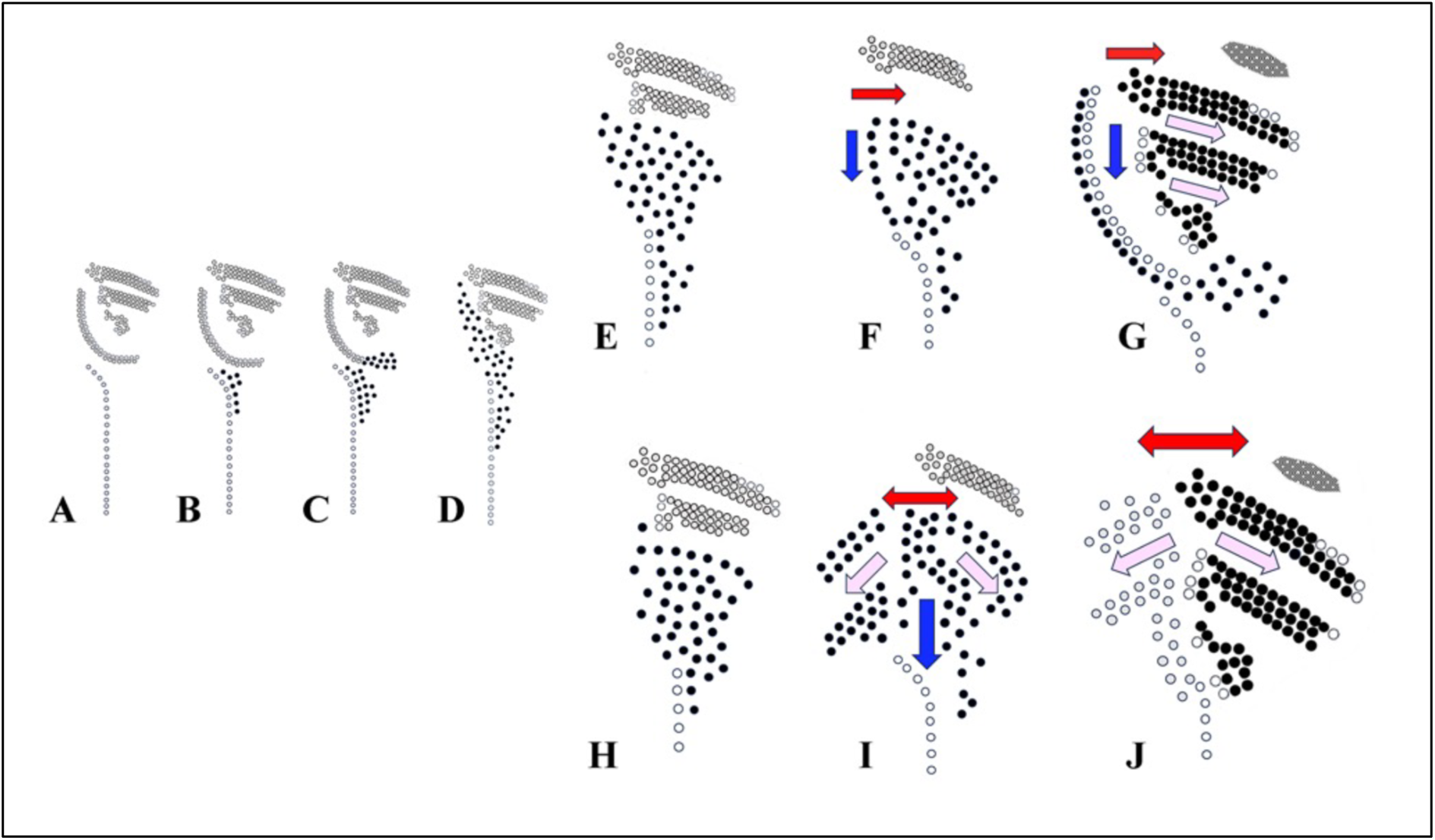
Oral replacement in a wildtype cell (A-G) and the ‘derangement’ of form in IA264 cells exposed to PLK inhibition using BI2536 (H-J). Parental basal bodies are depicted with open circles, while newly formed ORP basal bodies active in OA assembly are depicted as black filled circles (except in J, where left and right basal bodies are differentially filled or open for contrast). The oral replacement primordium (ORP) first appears as a field of basal body proliferation at the anterior end of the post-oral ciliary row (B). Shortly after, disassembly of the UM begins from the posterior end, and its basal bodies contribute to the proliferating field (C). The UM completely disassembles (D) and the ORP continues to replicate a field of basal bodies. One by one the primary membranelles (M1-M3) are either disassembled, or ultimately resorbed (E-G) (black arrow in figure G). As the parental membranelles are disassembled and resorbed, the ORP begins to form discrete, new membranelles (F,G). Membranelles assemble with distinctive asymmetry revealing organization along both the A/P axis and across the left/right field of symmetry. Our model suggests hypothetical ‘influences’ driving A/P organization within the developing membranelle field (Fig. F. turquoise arrow) and left/right organization (red arrow) resulting in a combined ‘vectorial’ influence (G. pink arrows) manifest in the diagonal orientation of the three primary membranelles and their anterior-right ‘sculpting’. We propose that inhibition of the cell’s PLK activity results in loss of polarity of the L/R ‘influence’, resulting in the symmetrical spread (double-red arrow, Fig. I). This results in a loss of UM assembly, and a mirrored set of primary membranelles (Figs. I, J).

In this model we suggest that PLK activity (in a wildtype/ control cell), is involved in driving local left/right asymmetry within the developing ORP (the oral replacement primordium). In the absence of PLK activity, this chiral-patterning influence spreads symmetrically both left and right of the UM and the post-oral ciliary row whose basal bodies contribute to the newly formed primordium. The result appears to be loss of the UM, an organelle that typically defines the left-most margin of the OA, and mirror-symmetric assembly of primary membranelles, M1-M3. In some sense, wildtype PLK activity appears to serve as a median firewall to an otherwise symmetric spread of OA-organizing activity, constraining its spread from the UM and post-oral meridian to the cell’s left.

When this firewall is abrogated by treatment with BI2536 (especially in cells homozygous for the *eja* mutation), this lateral patterning-influence spreads symmetrically, creating a mirrored membranelle pattern or ‘chevron’ configuration. An open question is whether the mirrored-membranelle pattern emerges during mid-body development of the oral primordium, or is somehow expressed only during rounds of oral replacement triggered by PLK inhibition.

### Thoughts on a symmetric, mirrored oral apparatus and the evolution of cortical pattern

On the face of it, it seems curious if not odd, that a loss-of-function mutation results, not simply in a disarranged ventral pattern of organelles, but in a mirror-duplication of the ventral pattern on the dorsal surface. This to our knowledge constitutes the only case of ‘intracellular homeosis’ ever described. Furthermore, no fewer than three separate genetic loci (*JANA, JAN-B,* and *JAN-C*) appear necessary to maintain this derived ‘singlet’ morphology. It is hard to escape the impression, that the default pattern for this ciliate: the ‘developmental ground-state’ to borrow language developed by Ed Lewis’ studies on pattern mutants in *Drosophila* (Lewis, 1951; Lewis, 1978; Gehring et al., 2009), is a cell in which oral structures form on both the dorsal and ventral hemi-cells, and that the dorsal organelles have been subsequently and robustly repressed.

It may be significant, that the most early-divergent ciliate species (some *Karyorelictid* species and most members of the class *Litostomatea* such as *Didinium*), exhibit symmetrical OAs, ones that circumscribe the entire anterior end of the cell (Fig 16). It is tempting to suppose that a symmetric, circumpolar OA represents the ancestral ciliate form, and that an asymmetric OA such as seen in *Tetrahymena*, and other *Hymenostome Oligohymenophorea*, is an evolutionarily derived form with members of the *Janus* gene family charged with suppressing dorsal stomatogenesis. This hypothesis is supported by the phylogenetic data that expose members of the *JANA* subclade of Polo kinases within the *Oligohymenophorea* but not within the more basal (early divergent) *Heterotrichea* (See Supplementary Fig 3).

**Figure 16.**
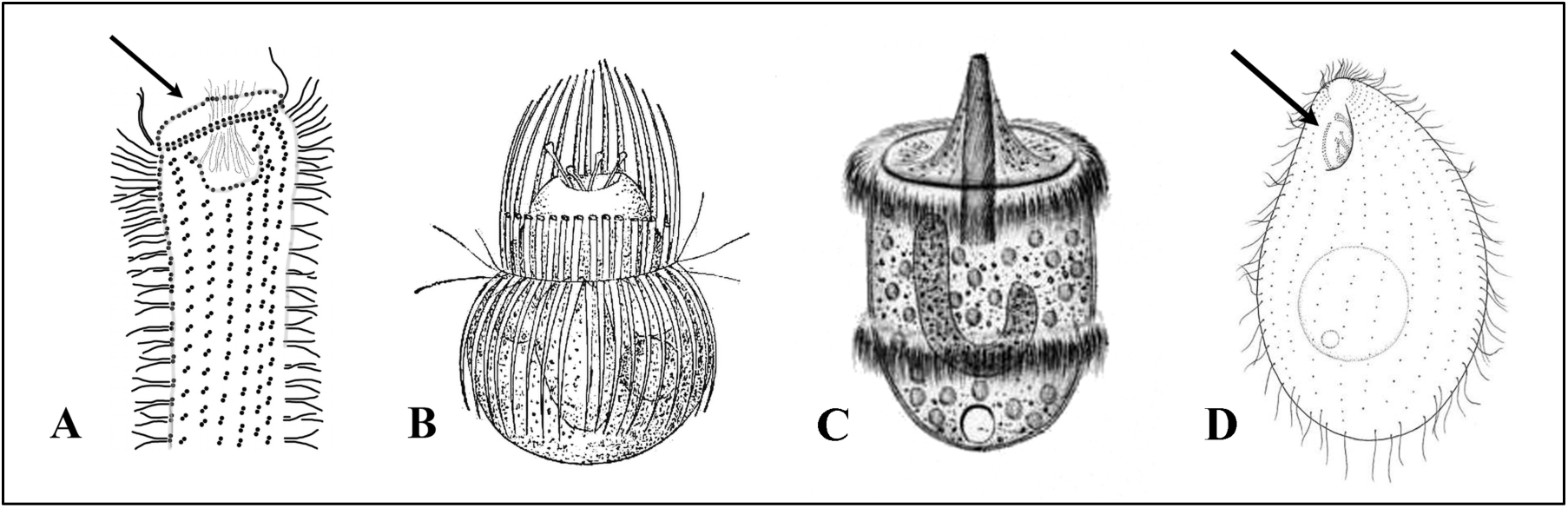
A) A Karyorelictid, an early divergent class of ciliate species showing a symmetrical OA that completely encircles the anterior aspect of the cell. (Modified from Foissner and Dragesco, 1996). B) *Mesodinium* another early divergent ciliate with a symmetrical, polar OA, (From Calkins, 1902). C) *Didinium*, a second ‘early divergent species from the class Litostomatea (From Stein, 1859). D) *Tetrahymena* exhibiting its asymmetric OA (from Lynn and Doerder, 2012).

### Cell division is linked to symmetry-breaking in both protozoan and metazoan cells

Despite their notoriety as cell-division / centriole replication-control factors, PLKs have also been implicated in non-cell-division functions, including symmetry breaking. Notable examples include polarization of the roundworm egg, C*aenorhabditis elegans* (Barbieri, et al., 2022; Kim & Griffin, 2021; Kim et al., 2024; Nishi, et al., and Velez-Aguilera, et al., 2024), and the asymmetric division associated with *Drosophila* neuroblast formation (Loyer & Januschke 2020; Noatynska, et al., 2013; 2008; Prehoda, 2009; Wodarz & Huttner 2003).

In the *Drosophila* neuroblast, as the centrosome replicates, the ‘dominant’ centrosome becomes asymmetrically localized to the apical cytoplasm (Rusan & Peifer, 2007; Januschke & Gonzalez, 2010). Polo (the original PLK) specifically localizes over this apical centrosome which organizes an interphase, mono-polarspindle. After neuroblast division, the cell inheriting the Polo-decorated centrosome maintains its stem-cell character. Polo (associated with the centrosome in the apical cortex) also phosphorylates the basal determinant Pon (‘partner of numb’) preventing it from docking in the apical cortex. PON therefore becomes concentrated in the basal cortex where it recruits Numb, a repressor of notch signaling (Wang, et al., 2007). Notch signaling subsequently controls stem cell proliferation in the apical daughter cell vs differentiation in the basal daughter cell. In this way, a centrosome-enriched kinase (Polo or PLK), associated with cell division exerts a role in driving intracellular asymmetry both by preferential decoration of a centrosome and by excluding proteins from a cortical domain.

Similarly, during lead-up to mitosis in the fertilized egg of *C. elegans*, the oocyte becomes polarized in a way that also involves a cell-cycle control Polo kinase, PLK-1. In this case, PLK-1 becomes physically anchored to the anterior cortex by association with anterior determinants and RNA-binding proteins Mex5/6. In a way similar to the *Drosophila* neuroblast, PLK-1 phosphorylates polarity determinant (POS-1) prohibiting it from docking at the anterior cortex and resulting in its concentration at the posterior cortex. POS-1 then plays a role in regulating gene expression in posterior cell descendants leading to germ-cell determination.

In both examples, a protein known for its roles in mitosis and control of cell-cycle events has been recruited to play a different role in driving intracellular patterning through establishment or maintenance of cellular polarity. It may be significant, that these activities play out within the cell cortex, defining cortical domains that recruit, or exclude other proteins and forming a specialized signal platform. It may also be significant that PLK is associated with the centrosome, a structure homologous to the basal body within ciliated or flagellated cells. Finally, PLK plays a chiefly inhibitory role, preventing the docking of cortical determinants so they come to occupy cortical territory in the hemi-cell opposite to where they are localized.

In *Tetrahymena*, we have identified a *PLK1* homolog *JANA*, that exhibits many of these same characteristics. The JanA product associates with basal bodies (similar to centrosomes) and is docked at the cell cortex. JanA is concentrated over basal bodies of the left-dorsal cell cortex where it appears to repress assembly of other organelle assembly platforms. Another PLK homolog (PLK2) appears to decorate basal bodies in the complementary, right-ventral hemi-cell (Fig 6E-H). What is novel, in this case, is that polar-localization of JanA occurs along the dorsal/ventral axis parallel to the axis of cell division, rather than the apical/basal or anterior/posterior axes perpendicular to the axis of cell division. Loss of JanA function generates a global, mirror-symmetric duplication of the entire ventral suite of cortical organelles in the dorsal *Tetrahymena* cortex, creating an intracellular ‘homeosis’.

## Methods

### Strains

Cell lines used in this study:

SD00630: IA220 janA-1/janA-1; eja1-1/eja1-1; chx1-1/chx1-1 (janA-1; eja1-1; chx1-1; janA, eja1, cy-r, II)

SD00631: IA221 *janA-1/janA-1; eja1-1/eja1-1; chx1-1/chx1-1* (*janA-1; eja1-1; chx1-1*; janA, eja1, cy-r, III)

SD00643: IA385 *janA-2/janA-2; eja1-1/eja1-1* (*janA-2; eja1-1*; janA, eja1, II)

SD00644: IA387 *janA-2/janA-2; eja1-1/eja1-1* (*janA-2; eja1-1*; janA, eja1, V)

SD00178: CU428 *mpr1-1/mpr1-1* (*MPR1*; mp-s, VII)

SD00632: IA264 *gal1-1/gal1-1; eja1-1/eja1-1* (*GAL1; eja1-1*; gal-s, eja1, II)

SD00015: A-star III

SD00022: B-star VI

*All strains provided by the Tetrahymena Stock Center now housed at Washington University, St. Louis.* https://sites.wustl.edu/tetrahymena/ *(formerly:* https://tetrahymena.vet.cornell.edu/).

### Cell Culture Conditions

Cells were grown at 30°C in ‘NEFF’ medium (0.25% proteose peptone, 0.25% dextrose, 0.5% yeast extract, 0.009% ferric EDTA). Matings were conducted using Dryl’s starvation medium medium (Dryl, 1959) (2 mM NaPO_4_ buffer, 2 mM sodium citrate, 1.5 mM CaCI_2_ pH 7.1).

### Identification of the janA-1 mutation

A mutant *T. thermophila* strain homozygous for *janA-1 and chx1-1* (IA220-RRID TSC_SD00630) was outcrossed to CU428-RRID TSC_ SD00178 (a 6-methyl purine-resistant heterokaryon), and the double drug-resistant (6-methylpurine [6mp] and cycloheximide [cy]) F1 progeny clones were selected. Several F1s were propagated vegetatively, and cy-sensitive macronuclear assortant clones were identified by replica plating. ‘Uniparental cytogamy’ (Cole, et al., 1992) was used to produce F2 progeny. To this end, a single cy-sensitive F1 was crossed to B*VI and the mating culture was exposed to a hyper-osmotic shock a 5.75 hrs into mating at 30°C. The shocked pairs were diluted and distributed to microtiter plates with Neff media to recover. Twenty-four hrs later, cycloheximide was added (*12.5 μg/ml)*, and cy-resistant survivors identified after four days in drug. 367 drug resistant synclones were identified. These were screened by eye for janus mutant appearance (slow-growing and abnormal shape), and screened again with high-resolution, DIC microscopy for the presence of secondary OAs. A total of 32 F2 clones with the *janA* phenotype and 41 clones with the wild-type phenotype were pooled. The pools were grown to a mid-log phase and starved for 2 d at room temperature in 60 mM Tris-HCl, pH 7.5. Total genomic DNA was extracted using the urea method (Dave *et al*., 2009). The pool DNAs were used to make genomic libraries using Illumina Truseq primer adapters and sequenced on an Illumina HiSeq X instrument, which generated paired-end reads of 150-base-pair length at 90× genome coverage. The MiModD suite of tools version 0.1.8 (https://sourceforge.net/projects/mimodd/) was used for ACCA-based variant mapping and identification (Jiang *et al*., 2017) as follows. The sequencing reads were aligned to the micronuclear reference genome (GenBank assembly accession GCA_000261185.1; Hamilton *et al*., 2016), and the aligned reads from both pools were used for joint multi-sample variant calling. The variant call dataset was filtered for sites with high coverage for each of the two pools. Linkage scores contrasting the allelic composition of the mutant with that of the WT pool were computed for each variant and the results plotted against micronuclear genome coordinates. For variant identification, the same sequencing reads were aligned to the macronuclear reference genome (GenBank assembly accession GCA_000189635 (Eisen *et al*., 2006) and the aligned reads subjected to variant calling as above.

Phylogenetic analyses were performed at the ngphylogeny.fr server (Lemoine, et al., 2019). Multiple sequence alignment was produced using MAFFT (Katoh and Standley 2013) and curated using trimAI (Capella-Gutierrez, et al., 2009). Neighbor-joining phylogenies were constructed using FastME and the statistical support of branches were calculated by bootstrap resampling with 1000 replicates Lefort, et al., 2015). Protein domains were detected using SMART (http://smart.embl-heidelberg.de/) tool and homologs in other species were identified using the Blastp tool at the Cildb database (http://cildb.i2bc.paris-saclay.fr/).

To knockout *TTHERM_00191790*, we prepared a plasmid for editing the *JANA* locus using homologous DNA recombination. To this end, we amplified two fragments of *TTHERM_00191790* using the primer pairs:

5’-CTATAGGGCGAATTGGAGCTGTTTGAGCTTACCTCAAGCCC-3’; CTAGAGCGGCCGCCACCGCCCACCTTTACCTAATAGCTTGCC-3’ and 5’-GCTTATCGATACCGTCGACCTGGTAGAGGATCAATGACTAACG-3’; 5’-AGGGAACAAAAGCTGGGTACAACTGTCTGGCTTTGCACCT-3’.

The fragments were cloned into the pNeo2_4 plasmid encoding the *neo5* selectable marker, to produce pNeo2_4_CDC5KO plasmid. The plasmid was linearized by digestion with PvuI and introduces into the wild-type strain CU428 or *eja1* homozygous strain IA264 by using biolistic bombardment (Cassidy-Hanley *et al*., 1997). Transformants were selected with 100 μg/ml paromomycin (pm). Transformant clones were grown in a progressively increasing concentration of pm to promote replacement of the endogenous locus by the disrupted locus due to phenotypic assortment.

To express a JanA-GFP fusion protein, we edited *TTHERM_00191790* by homologous DNA recombination using a plasmid pCDC5-GFP-3’5’, made by amplifying a fragment of the coding region with primers:

5’-CTATAGGGCGAATTGGAGCTAAGGAAATGCTCTCAAGCTTAATC-3’ and 5’-ATCAAGCTTGCCATCCGCGGGTTATAATTTTCCTTTTCTGTACCA-3’ and a fragment of the 3’ UTR with primers

5’-GCTTATCGATACCGTCGACCCTGAAGATTTAGGTCCCTCCC-3’ and 5’-AGGGAACAAAAGCTGGGTACGCATTTCTTTTTGGATCAGCTG-3’ and cloning into pGFP_neo2-4. pCDC5-GFP-3’5’ was linearized with SacI and KpnI and used for biolistic transformation of CU428 as described above.

To overproduce JanA, we inserted the MTT1-GFP sequence at the 5’ end of the coding region of *TTHERM_00191790*. The needed targeting plasmid, pMTT1-GFP-CDC5, was made by amplifying a portion of the 5’ UTR of *TTHERM_00191790* with primers 5’CTATAGGGCGAATTGGAGCTGCTTAGATAAATAGATAGATAGATAGTTAGTAAGTT-3’ and 5’-CTAGAGCGGCCGCCACCGCGTTGTTGAAGCTGAAGGGTTTC-3’ and a portion of the coding region with primers: TGAACTATACAAACGCGTCATGGATAAGGCAAACGAAG and AGGGAACAAAAGCTGGGTACCGGGCCTTGCTATTGAAATGAATACTCGCTAAG.

These fragments were subcloned into the pLF4gOE_neo2_4 plasmid in place of the fragments used to edit the *LF4A* gene (Jiang, et al. 2019b). pMTT1-GFP-CDC5 was linearized with PvuI, introduced by biolistic bombardment and transformants were selected with pm as described above. To induce overexpression, cells carrying the MTT1-driven gene copies were treated with cadmium chloride (2.5 mg/ml) in the standard growth medium SPPA.

### Immunofluorescence imaging

Cells were fixed and stained with antibodies as described by (Jiang et al., 2020). Samples were air-dried at 30°C, the cover glass was washed three times with PBS and incubated with primary antibodies diluted in PBS supplemented with 3% BSA fraction V and 0.01% Tween-20. The primary antibodies used were: polyclonal anti-GFP (Rockland; 1:800 dilution), monoclonal anti-centrin 20H5 (EMD Millipore; 1:400; Salisbury et al., 1988), and monoclonal anti-fenestrin (3A7) (Nelsen et al., 1994). The latter is now available from the DSHB located at the University of Iowa. Secondary antibodies were conjugated to either TRITC or FITC (Rockland; 1:300). The nuclei were co-stained with DAPI (Sigma-Aldrich). The labeled cells were embedded in VectaShield anti-quenching mounting medium for fluorescence (VectorLabs).

### Microscopy

Cells were imaged using an Olympus BX50 fluorescence microscope and a C-Mos digital camera. SR-SIM imaging was done on an ELYRA S1 microscope equipped with a 63× NA 1.4 Oil Plan-Apochromat DIC objective. Images were processed using ZEN 2011 and Fiji/ImageJ software.

### Drug treatment

Volasertib, (BI6727) was obtained from Sigma-Aldrich and dissolved in DMSO creating a 1mM concentrate. 10mM BI2536 in DMSO was obtained from ThermoScientific. We diluted it to a 1mM solution to create a stock solution. These stock solutions were then diluted into cell cultures to create the range of final concentrations.

## Acknowledgments

Research in the Cole laboratory was supported by NSF #1947608. Research in Gaertig lab was supported by the NIH grant 5R01GM135444. We appreciate the assistance of Liana R.X. Cole, in helping us to discern the Janus form from the morphologically similar, parabiotic-doublet form, a skill that proved essential in screening our initial pools of *janA* clones, and Kathleen R. Stuart for assistance in screening monoclonal antibodies for the fenestrin antigen. This article is dedicated to the years of extraordinary investigative work conducted in the laboratory of Professor Joseph Frankel, University of Iowa.

## Abreviations

CVP: contractile vacuole pore
Cyp: cytoproct
DIC: differential interference microscopy
eja: enhancer of janA
FZ: fission zone
GFP: green fluorescence protein
*JANA*: Janus A gene
JanA: janus A protein
*janA*: janus A mutant
MAC: macronucleus
MIC: micronucleus
MTT1: metallothionein 1
nCVP: new CVPs
OA: oral apparatus
OP: oral primordium
OR: oral replacement
PLK: polo-like kinase
pOM: primary oral primordium
SIM: structured illumination microscopy
sOP: secondary oral primordium
SPPA: growth medium: sugar, proteose peptone, antibiotics
UM: undulating membrane

