## Supplementary Figures for "The ‘Janus A’ gene encodes a polo-kinase whose loss creates a dorsal/ventral intracellular homeosis in the ciliate, *Tetrahymena*"

### Supplementary Materials

**A)**

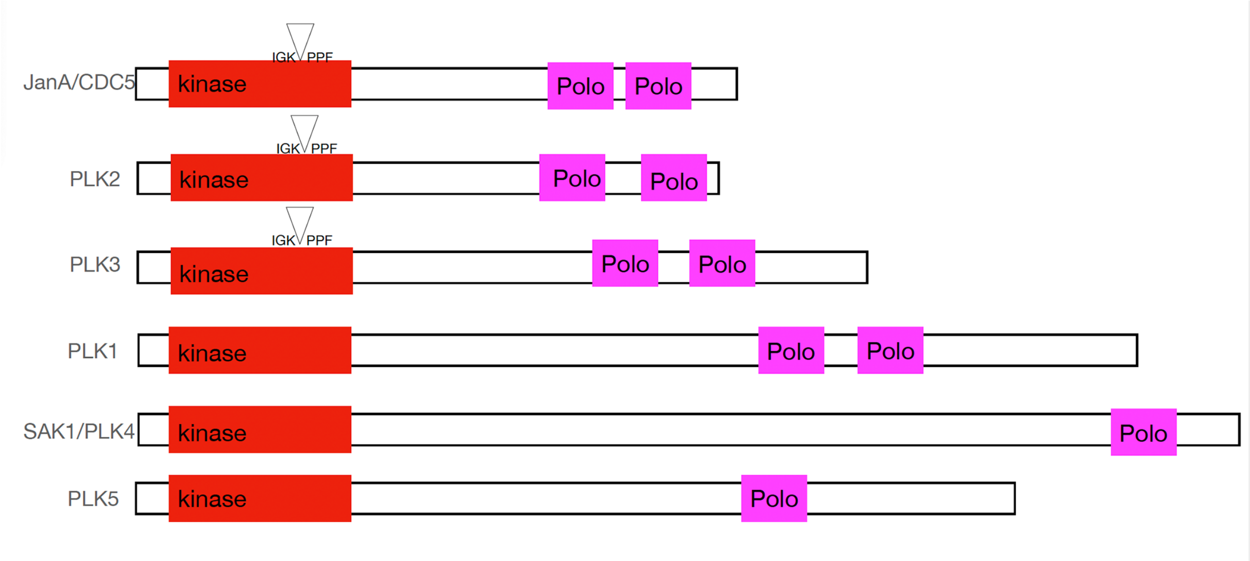


**B) TTHERM_00191790 (protein)**

MDKANEGKDQLIIIEEKISKVNGEMAVKKYLRGKLLGKGGFAKCYEVTNLETKKIIAAKI
IPKQTLTKNRARQKLISEIKIHKSLNHQHVVAFEHVFEDHENVYILLELCTNHTLNELIK
RRKRLTELEVQCYVVQIVNALKYLHSHKVIHRDLKLGNLFLNEKMEIKLGDFGLATKLEF
DGEKKHTICGTPNYIAPEILDGKTGHSYQVDIWSLGVIIYTLLIGKPPFETPDVKTTYKK
IRNISYGFPENVPISDQARGLITRILNIDPQRRPTLDEIMSSSFLNTGGTIPKVLPLATL
ACPPSASYTKQFLPQGNALKLNQAPMRLTDSASQSTTNIKGKNASTSNLYSKQTTTGAVP
DASRPLGGTTSNWNQVSSQKDQRPQTQQFQKSDFKTSMSTKNMGSGSQNPLFNQTNAPGL
ATAQSLNTFTKTKNNFNSTQKGFMQTGTQNFGKPVQPVWVTQWVDYSAKYGLGYLLSNGC
SGVFFNDSTKVILDPTTNNIEYIERLGNERQDFVQQFTLKDYPPEMKKKVTLLSHFKSYL
EEQMQKQNIQIEVPQFQGDLQPYVYVKKWMKTKHAIMFRLSNKIVQVSFQDKTEILLSSE
NKMVTYVDRKGTRTDYPLSTALESSNTEMSKRLKYTKDILTHMLNNNTTNNNNGRGSMTN
VNPTQNGTEKENYN

**Supplementary Figure 1.** A) The protein domain organization of Polo kinases of *Tetrahymena*. The triangles mark the position of an intron in the corresponding coding regions of genes, that is conserved in *JANA*, *PLK2* and *PLK3*, arguing that these three Polo kinases are paralogs.
B) The predicted JanA gene product. Turquoise = catalytic domain of a PLK: Serine/Threonine kinase. {Residues 28-285}. E-value: 2.15 $e^{-168}$. Green = first Polo-box. {Residues: 456-546). E-value: 1.85 $e^{-35}$. Yellow = second Polo-box. {Residues: 564-643}. E-value: 3.56 $e^{-30}$. Sites of *janA* mutations are indicated in magenta. The *janA-1* allele had gained a STOP codon at residue W213. The *janA-2* mutant allele exhibits a frameshift starting at K185.

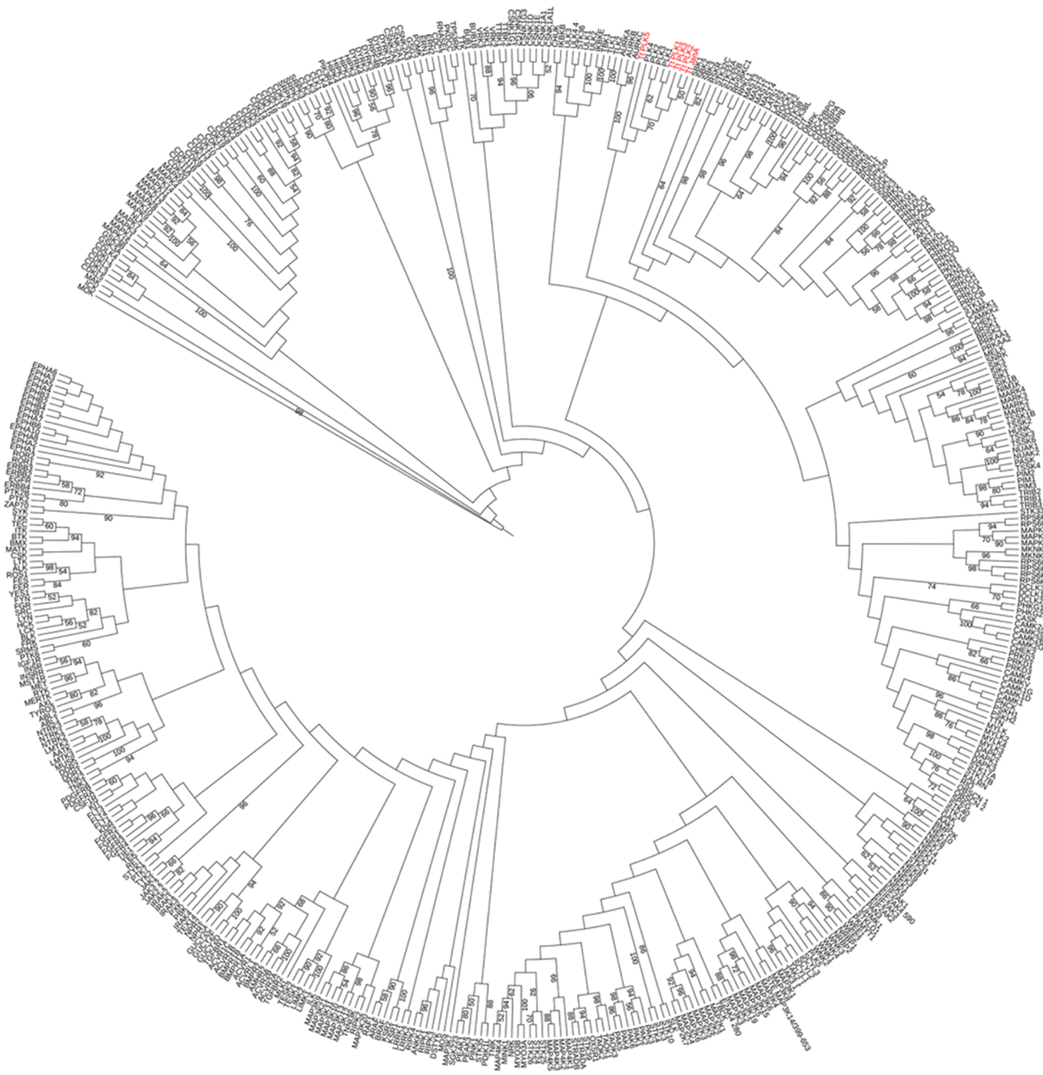

**Supplementary Figure 2.** A neighbor-joining phylogenetic analysis performed using 497 sequences of human kinase domains (Modi and Dunbrack, 2019b) and the corresponding sequences of *Tetrahymena JANA* and four closest homologs based on BlastP searcher. The numbers represent statistical support for branches based on 1000 bootstrap resamples. The tree is rooted to the human PTK7. The five *Tetrahymena* sequences are labeled in red.

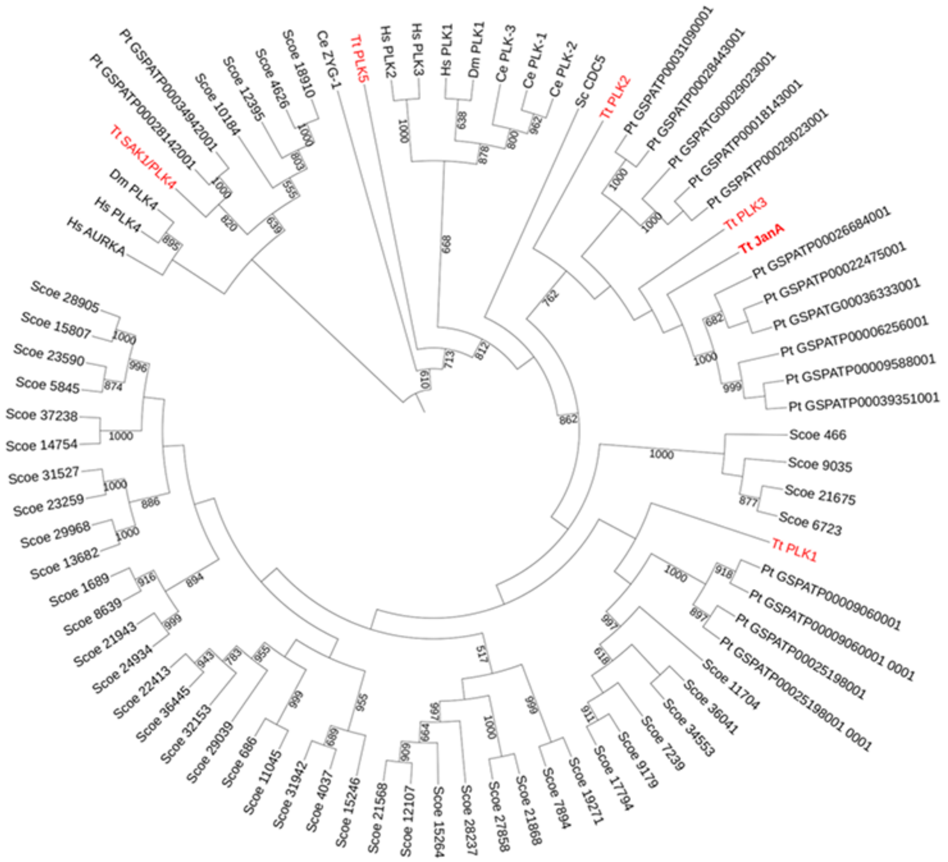


**Supplementary Figure 3.** A neighbor-joining phylogenetic analysis of Polo kinase activity in ciliates, and other eukaryotes. The sequences in ciliate genomes (*Tetrahymena* were identified by BlastP searches using established Polo kinase sequences such as *CDC5*) and potential orthologs were selected based on the high homology within the kinase activity domain and the presence of POLO box domains. Note that all ciliate proteins belong to a clade with animal *PLK1*, *PLK2* and *PLK3* and *CDC5* of *S. cerevisiae*. *JANA* and *Tetrahymena* paralogs *PLK2* and *PLK3* belong to a subclade with members present in *Paramecium tetraurelia* but not in *Stentor coeruleus*.

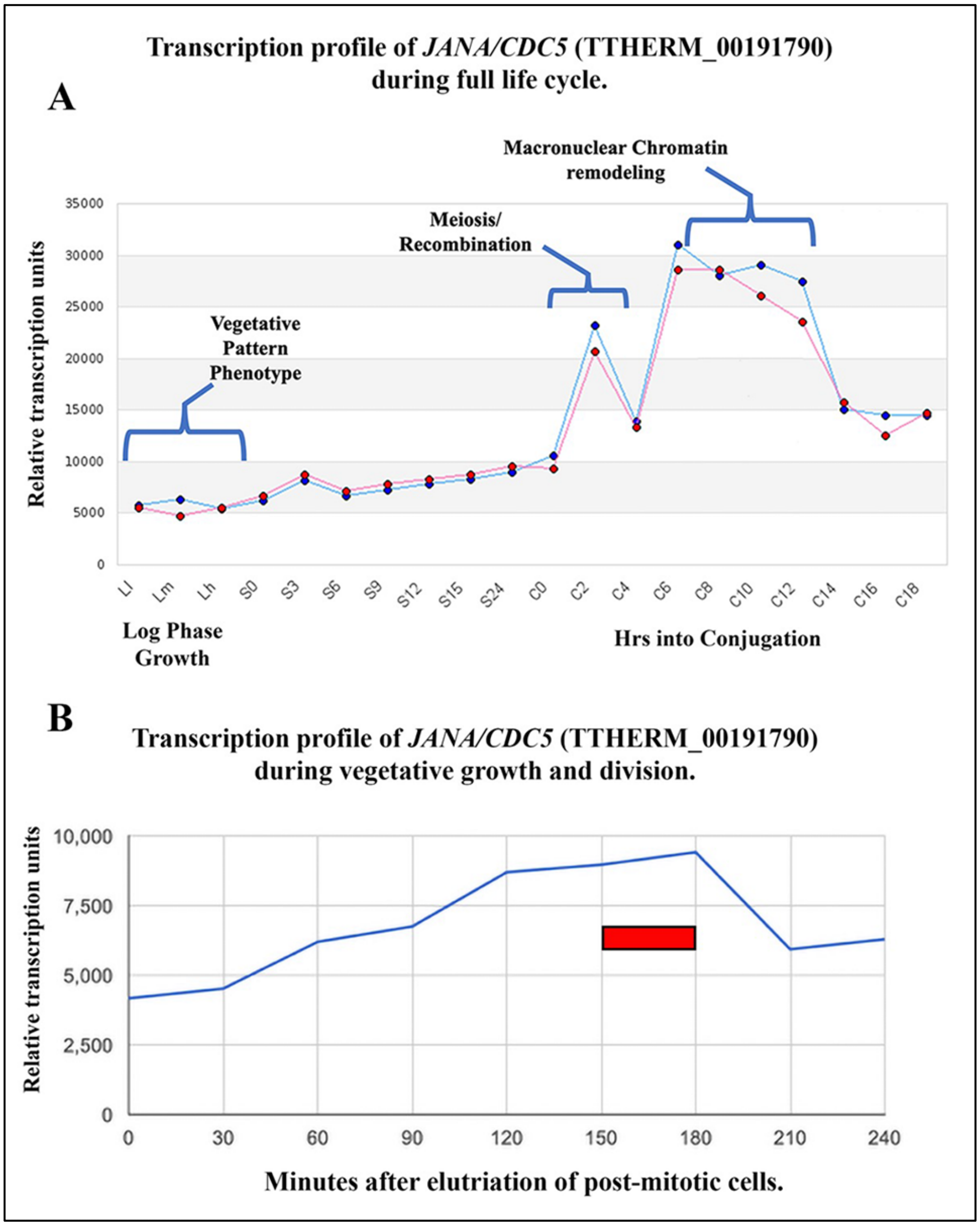


**Supplementary Figure 4**. Expression profile of mRNA for *JANA*/*CDC5* (TTHERM_00191790) obtained from the Tetrahymena Functional Genomics database (http://tfgd.ihb.ac.cn/search/detail/gene/TTHERM_00191790). A) The levels of mRNA during various life-history stages: L-l, L-m and L-h: vegetatively growing cells collected at ~1x10$^5$ cells/ml, ~3.5x10$^5$cells/ml and ~1x10$^6$ cells/ml. S-0, S-3, S-6, S-9, S-12, S-15 and S-24: cells starved for 0, 3, 6, 9, 12, 15 and 24 hours. C-0, C-2, C-4, C-6, C-8, C-10, C-12, C-14, C-16 and C-18: conjugating cells collected at 0, 2, 4, 6, 8, 10, 12, 14, 16 and 18 hours after initiation of conjugation by mixing different mating types. B) *JANA*/*CDC5* (TTHERM_00191790) mRNA levels during vegetative growth (minutes after centrifugal elutriation enriched a population of post mitotic cells). Red bar indicates the approximate time of cytokinesis.
